# A new approach to the determination of tubular membrane capacitance: passive membrane electrical properties under reduced electrical conductivity of the extracellular solution

**DOI:** 10.1101/2021.11.12.468264

**Authors:** Jiří Šimurda, Milena Šimurdová, Olga Švecová, Markéta Bébarová

## Abstract

The tubular system of cardiomyocytes plays a key role in excitation-contraction coupling. To determine the area of the tubular membrane in relation to the area of the surface membrane, indirect measurements through the determination of membrane capacitances by electrophysiological measurements are currently used in addition to microscopic methods. Unlike existing electrophysiological methods based on an irreversible procedure (osmotic shock), the proposed approach uses a reversible short-term intermittent increase in the electrical resistance of the extracellular medium. The resulting increase in the lumen resistance of the tubular system makes it possible to determine separately capacitances of the tubular and surface membranes from altered capacitive current responses to subthreshold voltage-clamped rectangular pulses. Based on the analysis of the time course of capacitive current, computational relations were derived which allow to quantify elements of the electrical equivalent circuit of the measured cardiomyocyte including both capacitances. The exposition to isotonic low-conductivity sucrose solution is reversible which is the main advantage of the proposed approach allowing repetitive measurements on the same cell under control and sucrose solutions. In addition, it might be possible to identify changes in both surface and tubular membrane capacitances caused by various interventions. Preliminary experiments in rat ventricular cardiomyocytes (*n* = 10) resulted in values of the surface and tubular capacitances 72.3 ± 16.4 and 42.1 ± 14.7 pF, respectively, implying the fraction of tubular capacitance/area of 0.36 ± 0.08. We conclude that the newly proposed method provides results comparable to those reported in literature and, in contrast to the currently used methods, enables repetitive evaluation of parameters describing the surface and tubular membranes. It may be used to study alterations of the tubular system resulting from various interventions including associated cardiac pathologies.

**Author summary:** The cell membrane of cardiomyocytes is invaginated forming a net of tubules through the whole cell (the tubular system). These invaginations are essential for the coordinated contraction of cardiomyocytes. Various cardiac diseases may lead to an alteration of the tubular system and *vice versa* alteration of the tubular system may result in cardiac dysfunction. Therefore, it is desirable to investigate the tubular membrane separately. Unfortunately, it is not easy and methods currently used for this purpose provide diverse results. The widely used detubulation methods (separating the tubular and surface membrane by an osmotic shock) are irreversible. It makes impossible repeated measurements on the same cell, thus, disabling evaluation of parameters describing the surface and tubular membranes and particularly testing changes in the tubular system. We developed an alternative approach based on the electrical separation of the tubular and surface membranes induced by exposition of the cardiomyocyte to an isotonic sucrose solution with low electrical conductivity. It allows to quantify the approximate area of the surface and tubular membrane using evaluation of electrical capacitance of both membrane areas. Measurements can be repeated several times on the same cell.

## Introduction

The membrane capacitance (*C*_m_) measured by electrophysiological methods can be considered a measure of the cell membrane area using the relationship

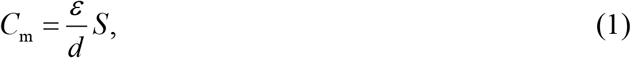

where *ε* is the permittivity, *d* the thickness, and *S* the area of the membrane. This applies provided the ratio *ε*/*d* (representing the specific membrane capacitance) is constant over the entire area of the membrane.

Measurements of the electrical parameters characterizing cellular membranes (separating the external and internal environment of cardiomyocytes) are complicated by complex membrane geometry, especially by the existence of the tubular system (t-system). Given the physiological importance of the t-system (see reviews [1, 2]), it is desirable to investigate the properties of the surface and tubular membranes separately. An important first step is to determine the area of the surface and tubular membrane.

If we consider *ε*/*d* to be a constant in the whole membrane system, the ratio of tubular and surface capacitance *k* = *C*_t_ / *C*_s_ equals the ratio of corresponding areas

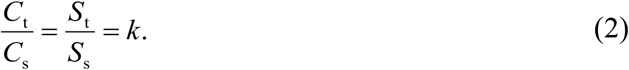

In the cardiomyocytes, simple electrophysiological measurements do not allow to assess both capacitances separately because the surface and the t-tubular systems are tightly electrically coupled. In terms of the model with lumped parameters (Fig 1B), the surface and tubular membranes are separated by the electrical resistance of the lumens of the t-tubular system which is so small that the responses of the membrane current to subthreshold steps of the applied membrane voltage (descending part of the capacitive current) follow a simple exponential waveform (for a detailed analysis see Appendix 2 of [3]). It follows that only the total membrane capacitance *C*_m_ = *C*_t_ + *C*_s_ of the whole membrane system can be estimated from usual electrophysiological experiments and parameters resulting from the mono-exponential approximation of descending part of the capacitive current.

**Fig 1.**
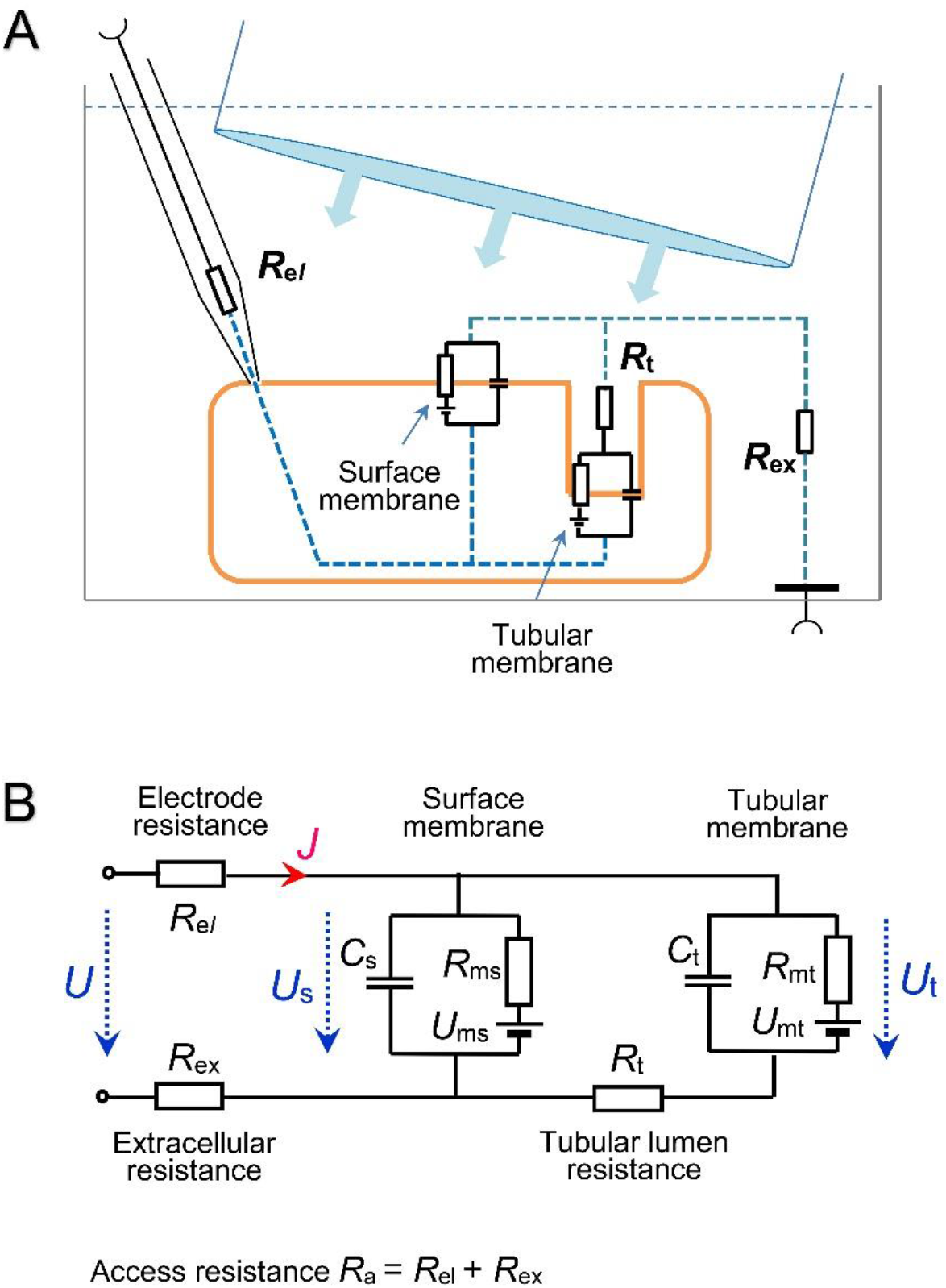
Principle of the method. (A) Experimental setup comprising an isolated cell, a glass microelectrode, and a jet pipe for rapid exchange of extracellular solutions. (B) Lumped-element electrical equivalent circuit of a cell with the developed tubular system connected to the measuring equipment. *R*_el_ – glass electrode resistance; *R*_ex_ –resistance of the extracellular solution between the electrode and the measured cell; *R*_a_ – access resistance (*R*_el_ + *R*_ex_); *C*_s_, *C*_t_ – membrane capacitances; *R*_ms_, *R*_mt_ – membrane resistances; *U*_ms_, *U*_mt_, – resting voltage of the surface and tubular membrane; *R*_t_ – resistance of the lumen of tubular system. *U, U*_s_, and *U*_t_ – imposed, surface, and tubular membrane voltage, respectively; *J* – membrane current.

To determine capacitances *C*_s_ and *C*_t_ electrophysiologically, the detubulation methods were developed consisting in electrical disconnection of tubular membranes by osmotic shock [4–7]. However, this widely used method is irreversible, which makes repeated measurements on the same cell and the use of paired difference tests impossible. In addition, the difficult-to-determine fraction of tubules may remain intact after the detubulation procedure [6, 8, 9] which may limit the accuracy of the *C*_t_ and *C*_s_ determination.

The basic idea of the proposed method is the electrical separation of the surface and tubular membrane system by increasing the electrical resistance of the t-tubule lumens. It can be expected that a marked reduction of the electrical coupling between the two membrane systems will transform the virtually mono-exponential course of the recorded capacitive current into two distinguishable exponential components, of which both membrane capacitances *C*_t_, *C*_s_, and the ratio of corresponding areas *k* (Eq (2)) could be calculated. An increase in the resistance separating the tubular system from the surface membrane can be achieved experimentally by a short-term transient replacement of the extracellular solution with the isotonic sucrose solution. This approach would leave the cell intact and allow repeated measurements.

This study is based on the analysis of the electrical equivalent circuit of a cardiomyocyte in a measuring arrangement. The electrical equivalent circuit with lumped parameters was used and described mathematically to derive formulas for the calculation of membrane capacitances *C*_t_, *C*_s_, and other elements of the equivalent circuit. The method was then verified using a computer model and tested in preliminary experiments.

## Results

### Theoretical background

Fig 1 illustrates a schematic representation of a simple electrical equivalent circuit with lumped parameters of cardiac cell connected to a glass microelectrode. In the subthreshold range of membrane voltage, the membrane resistances are regarded as constants and the measured system is described mathematically by a system of two non-homogeneous linear differential equations of the first order with respect to time (*t*) for variables *U*_s_ and *U*_t_ representing surface and tubular membrane voltages.

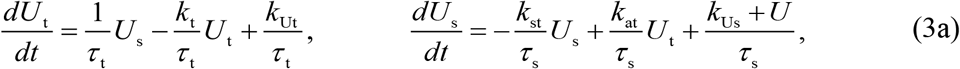

where

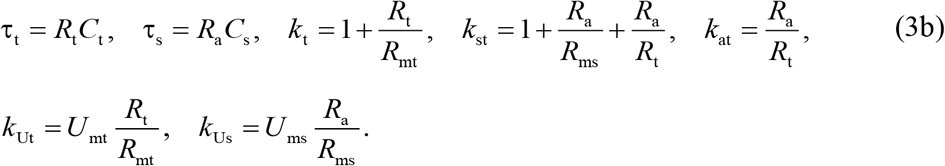

The Eq (3a,b) can be solved following the standard approach to systems of linear non-homogenous differential equations (e.g. [10]). In the particular case of the response to the imposed subthreshold step of membrane voltage *U* from the level *U*_1_ to *U*_2_, the solution of Eq (3a,b) leads to a sum of two exponential functions

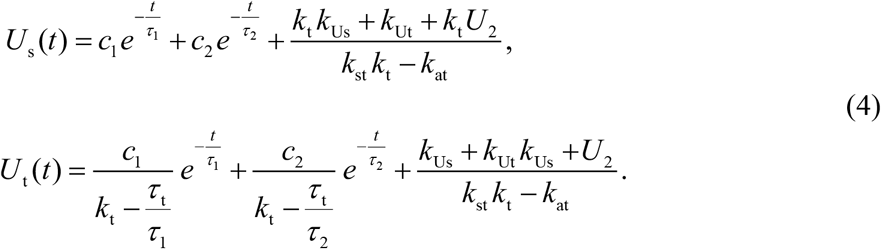

Considering initial conditions, the constants *c*_1_ and *c*_2_ are expressed as

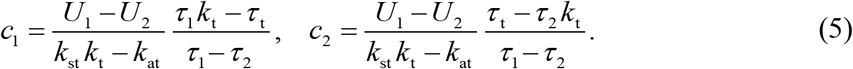

The time constants *τ*_1_, *τ*_2_ of the two exponential terms satisfy the conditions arising from the properties of the roots of the characteristic equation

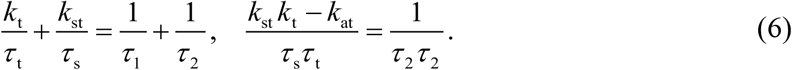

The only measured quantity is membrane current *J*, which is simply related to membrane voltage *U*_s_ by the Ohm law

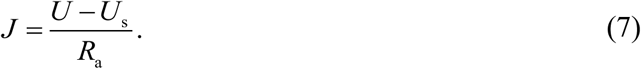

Considering the Eqs (4) and (7), the response of the current *J* to a small step of membrane voltage (from *U*_1_ to *U*_2_) can be expressed as a sum of two exponential terms and a constant

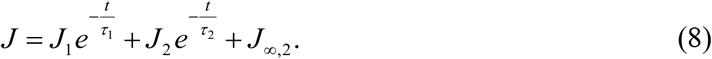

*J*_∞,1_ and *J*_∞,2_ denote steady-state currents at the membrane voltages *U*_1_ and *U*_2_. The bi-exponential approximation of the recorded current provides numeric values of six parameters, namely *J*_1_, *J*_2_, *J*_∞,1_, *J*_∞,2_, *τ*_1_, and *τ*_2_. Their relation to the parameters of the model (Eq (3b)) is given by a set of four Eqs (9)

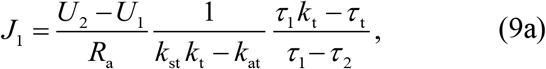

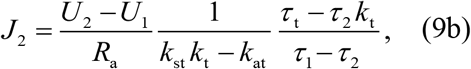

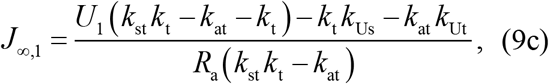

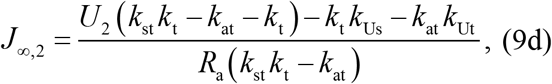

supplemented by two Eqs (6).

Only a limited number of the elements of the equivalent circuit in Fig 1B can be determined from the six parameters *J*_1_, *J*_2_, *J*_∞,1_, *J*_∞,2_, *τ*_1_, and *τ*_2_ estimated from the recorded capacitive current.

### Elements of the electrical equivalent circuit

The access resistance *R*_a_, the time constant *τ*_s_, and the capacitance of the surface membrane *C*_s_ could be expressed from Eq (3b), (6), and (9) after rearrangements:

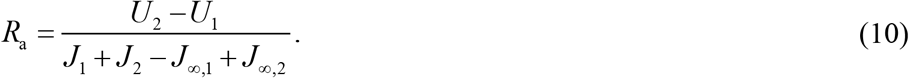

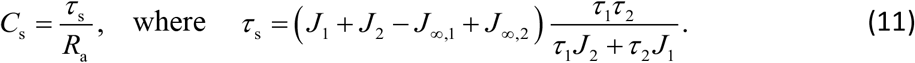

The resistances of tubular membrane and tubular lumen could not be expressed directly from Eq (3b), (6), and (9). However, two combinations of resistances *R*_ms_, *R*_mt_, and *R*_t_ (denoted *R*_1_ and *R*_2_) could be calculated as

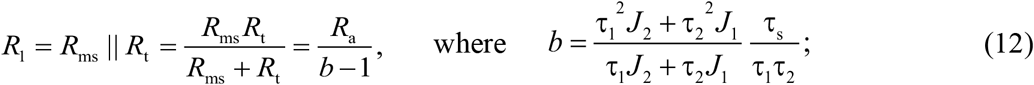

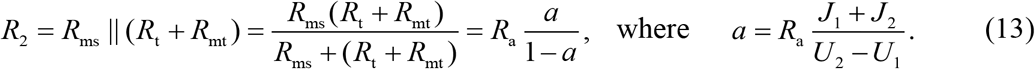

For convenience, parallel combinations of resistances were expressed by the symbol ||. This notation will be retained in the whole text.

The tubular membrane capacitance *C*_t_ could theoretically be expressed as

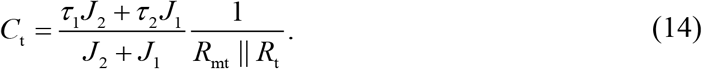

However, Eq (9) and (6) did not allow to express formula for the calculation of parallel combination *R*_mt_ || *R*_t_. As mentioned in the Discussion, we looked at two ways to solve this problem. In this work, we prefer the introduction of an additional assumption. It is reasonable to expect the ratio of membrane conductance *G*_mt_/*G*_ms_ (= *R*_ms_/*R*_mt_) to be proportional to the ratio of membrane areas like the ratio *C*_t_/*C*_s_ according to Eq (2).

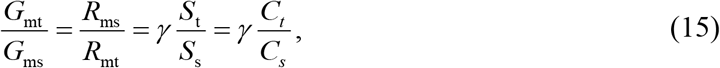

where *ɣ* is a hitherto unknown coefficient of proportionality, the value of which may be different from 1 regarding the heterogeneity of tubular membrane (namely due to the differences in the distribution of ionic channels).

To determine the value of *k* from Eq (2) and (15), we used definitions of the resistances *R*_1_ and *R*_2_ (directly computable from Eq (12) and (13)). After introducing the ratio *R*_1_/*R*_2_ into (15) and adjusting, we arrived at a quadratic equation for *R*_ms_ as a function of *ɣk*

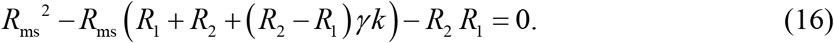

Only one root of the Eq (16)

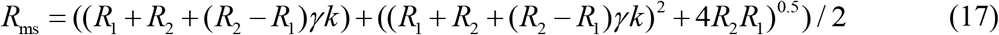

led to a physically realistic solution.

By inserting expressions of *C*_t_ and *C*_s_ (Eqs (14) and (11)) into Eq (2), and considering the definition of *R*_1_ (Eq (12)), we got another expression of *R*_ms_ as a function of *k* and *ɣ*

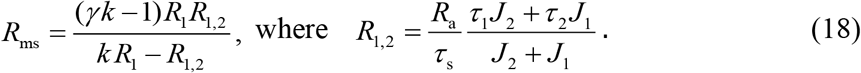

The numeric values of the variables *R*_ms_ and *k* could be calculated (for the selected *ɣ* value) from the system of two equations (Eqs (17) and (18)). The constant *k* could also be calculated from one implicit equation after comparing the right sides of the Eqs (17) and (18).

The next section will show how the calculated value of *k* and the membrane capacitances (*C*_t_, *C*_s_) depend on the *ɣ* setting. All the constants in the Eq (17) and (18) can be calculated from the parameters *J*_1_, *J*_2_, *J*_∞,1_, *J*_∞,2_, *τ*_1_, and *τ*_2_ determined from the results of fitting the membrane current response (to a small voltage step) by the sum of two exponential functions and a constant. Calculation of the constant *k* makes it possible to quantify other elements of the electrical equivalent circuit (Fig 1B). In addition to the expressions derived so far for *R*_a_, *C*_s_, and *R*_ms_ (Eqs (10), (11), and (18)), the remaining elements can be easily calculated. The most important parameter *C*_t_ follows from Eq (2)

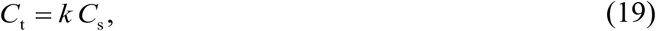

the resistances *R*_t_ and *R*_mt_ from Eq (12) and (15) as

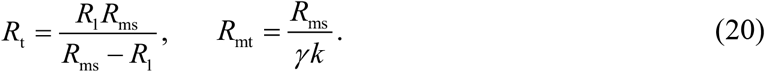

The total membrane capacitance and the fraction of tubular capacitance can be expressed as

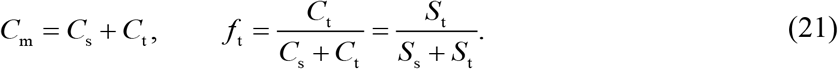

The formulas allowing quantification of the elements of the electrical equivalent circuit are summarized in Table 1.

**Table 1.**
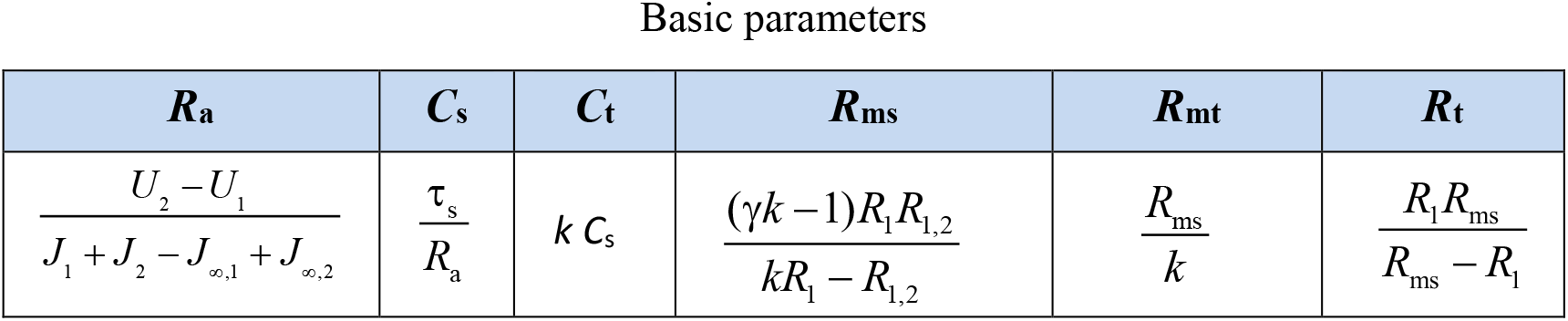
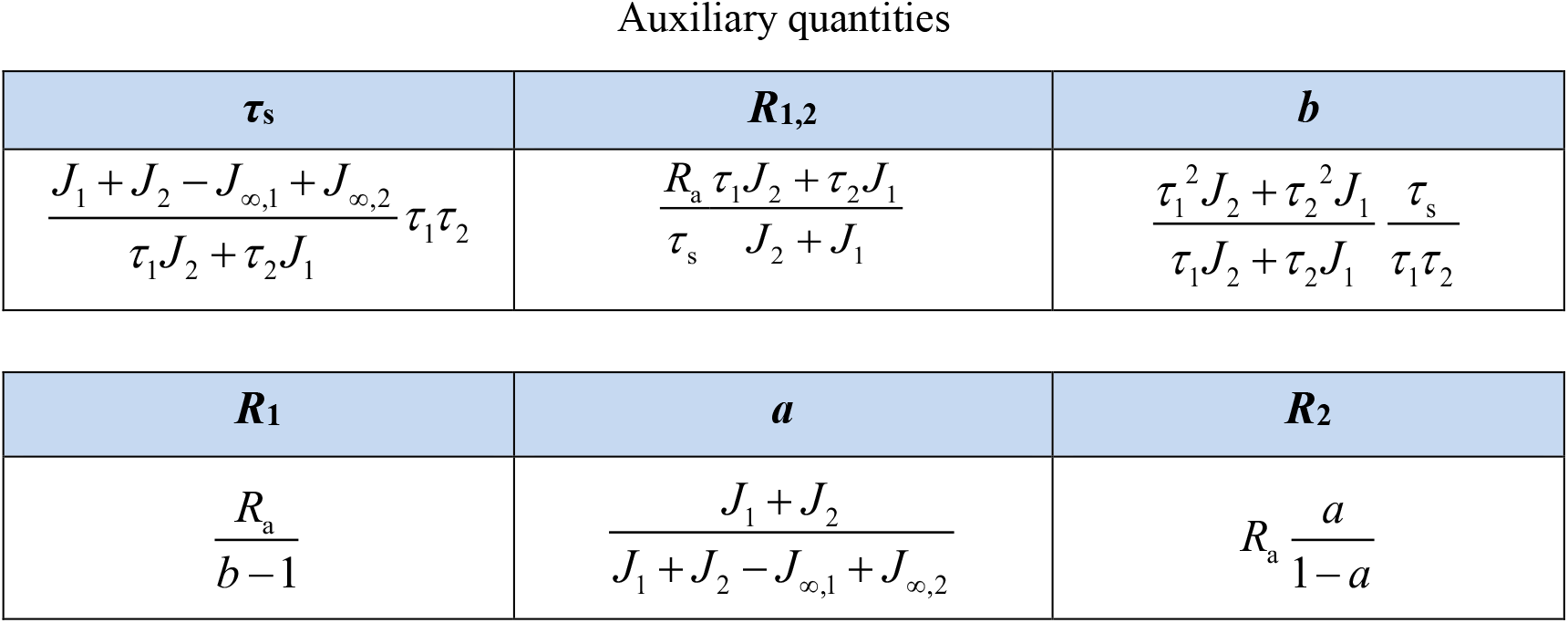
Mathematical formulas for calculation of the elements of electrical equivalent circuit. Calculation of the elements of electrical equivalent circuit (Fig 1B) from the parameters *J*_1_, *J*_2_, *J*_∞,1_, *J*_∞,2_, *τ*_1_ and *τ*_2_ resulted from a double-exponential analysis of current response to a subthreshold step of membrane voltage. The value of *k* results from the solution of the Eqs (17) and (18).

The values of the reversal voltages *U*_ms_ and *U*_mt_ can be estimated from the parameters *J*_1_, *J*_2_, *J*_∞,1_, *J*_∞,2_, *τ*_1_, and *τ*_2_ only approximately under the assumption that *U*_ms_ = *U*_mt_ (which may not be exactly met): the relations *U*_ms_ = *U*_mt_ = *U*_1_ – *J*_∞,1_ *R*_a_ /(1-*a*) = *U*_2_ – *J*_∞,2_ *R*_a_/(1-*a*) follow from Eq (7c,d). However, the calculated values of *C*_t_, *C*_s_, and *f*_t_ are independent of the values of *U*_ms_ and *U*_mt_ used for calculations.

### Model verification of the theory

To proof the correctness of the described calculations of the elements of the electrical equivalent circuit, we proposed a software written in MATLAB Live Editor (S1_verification.mlx - available as Supporting information) based on the solution of the set of differential equations (Eqs (3a,b)). The software was designed to mimic real experiments on isolated cells. The numerical values of *R*_a_, *R*_t_, *R*_ms_, *R*_mt_, *C*_s_, *C*_t_, *U*_ms_, and *U*_mt_ are optional. The values summarized in Table 2 were chosen as examples of values close to those obtained from preliminary experiments. The voltage *U*_2_ was set to −75 mV, however, this option did not affect the calculated values of the elements of the electrical equivalent circuit. The results obtained by the calculations are compared in the following figures with the real values set according to Table 2.

**Table 2.** Values of the elements of electrical equivalent circuit used for verification of the derived formulas.

| $R_a$ [MΩ] | $R_t$ [MΩ] | $R_{ms}$ [MΩ] | $R_{mt}$ [MΩ] | $C_s$ [pF] | $C_t$ [pF] | $U_{ms}$ [mV] | $U_{mt}$ [mV] |
| --- | --- | --- | --- | --- | --- | --- | --- |
| 12.5 | 15 | 150 | 241 | 74 | 46 | -160 | -160 |

The shape of the imposed rectangular voltage impulse mimicking experimental records with limited rising and falling edge (Fig 2A) resulted from simultaneously solved additional simple differential equation to create fast exponential onset and offset of the imposed impulses with the optional time constant (*τ*_p_ = 0.05 ms was used in most computations). Fig 2B shows computed responses of surface and tubular membrane voltage (*U*_s_ and *U*_t_). The characteristics of the experimental capacitive current with a steep increase to a maximum followed by a slow decay are reproduced in the simulated current response (Fig 2C).

**Fig 2.**
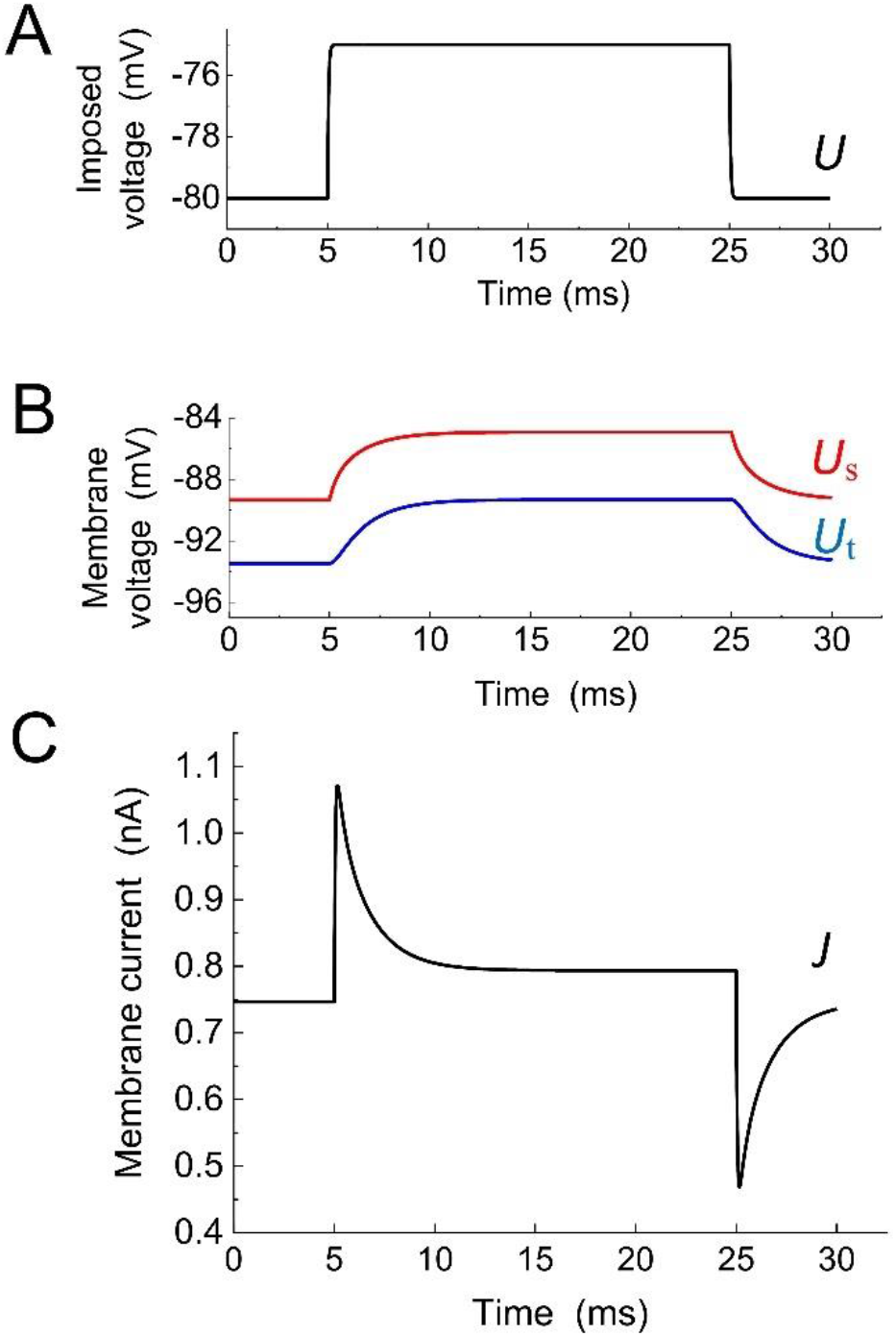
Solution of Eq (3a,b) describing the electrical equivalent circuit shown in Fig 1B. Selected parameter values are given in Table 2. (A) Imposed voltage impulse *U*; an additional simple differential equation was simultaneously solved to create slower rising and falling edge of the rectangular voltage impulse closer to actual shape. (B) Responses of surface and tubular membrane voltage *U*_s_ and *U*_t_. (C) Response of the membrane current *J*.

The descending phase of the simulated capacitive current was expected to follow a distinct bi-exponential course because the tubular resistance *R*_t_ was set to a sufficiently high value corresponding to the effect of sucrose solution. Using the *Curve fitting tool* of MATLAB (R 2017a), a section of capacitive current marked in Fig 3 was fitted to the bi-exponential function expressed by Eq (8) as in evaluation of experimental results. The resulting numeric values of five parameters *J*_1_, *J*_2_, *J*_∞,2_, *τ*_1_, and *τ*_2_ supplemented by the resting current *J∞*,_1_ were subsequently inserted into Eqs (10)–(13), (17)–(21) to calculate the elements of electrical equivalent scheme.

**Fig 3.**
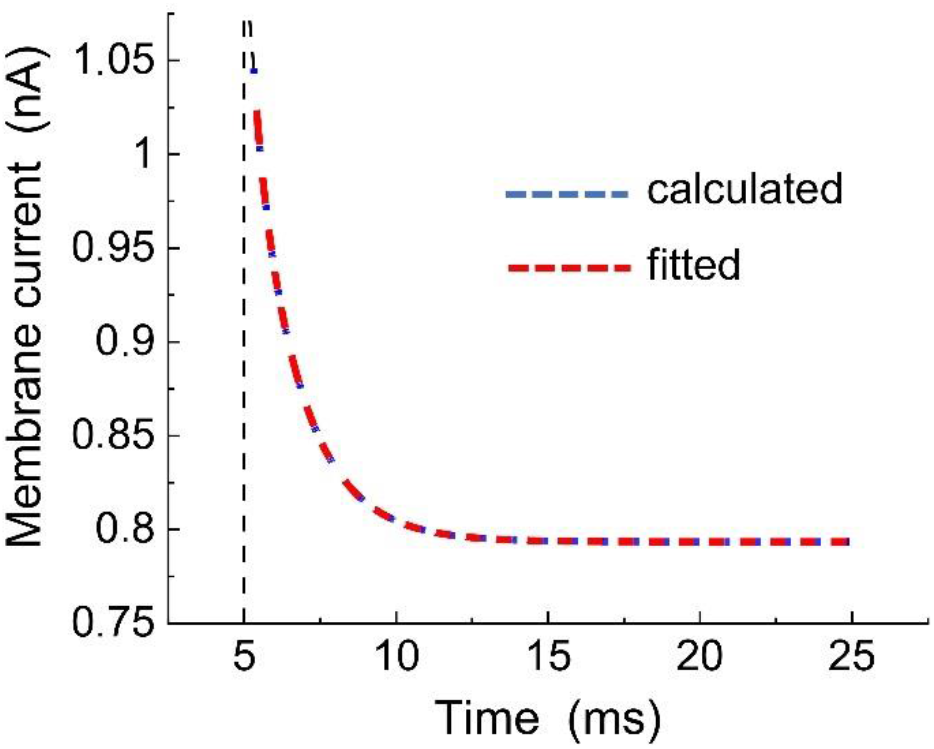
Approximation of the capacitive current by the sum of two exponential functions and a constant. The evaluation started 0.4 ms after the onset of the depolarization step. The part of capacitive current (shown in Fig 2) was calculated as a solution of differential Eq (3a,b) with values according to Table 2 (blue) and fitted with bi-exponential function using the *Curve fitting tool* in Matlab (red) resulting in *J*_1_ = 0.2677 nA, *J*_2_ = 0.0895 nA, *J*_∞,2_ = 0.7104 nA, *τ*_1_ = 1.6285 ms, and *τ*_2_ = 0.3382 ms.

The key point was the estimation of the surface membrane conductance *G*_ms_ = 1/*R*_ms_ and the constant of proportionality *k* between the tubular and surface membrane capacitances. The value of the constant *ɣ* related to the ratio of membrane conductance was still unknown. To verify the derived formulas, we first solved the system of two equations (17) and (18) assuming *ɣ=G*_mt_*C*_s_/(*G*_ms_*C*_t_) to meet exactly Eq (15). The choice of parameters according to Table 2 resulted in *ɣ*~1. The numerical values of *G*_ms_ and *k* corresponded to intersection of the two plotted curves as illustrated in Fig 4.

**Fig 4.**
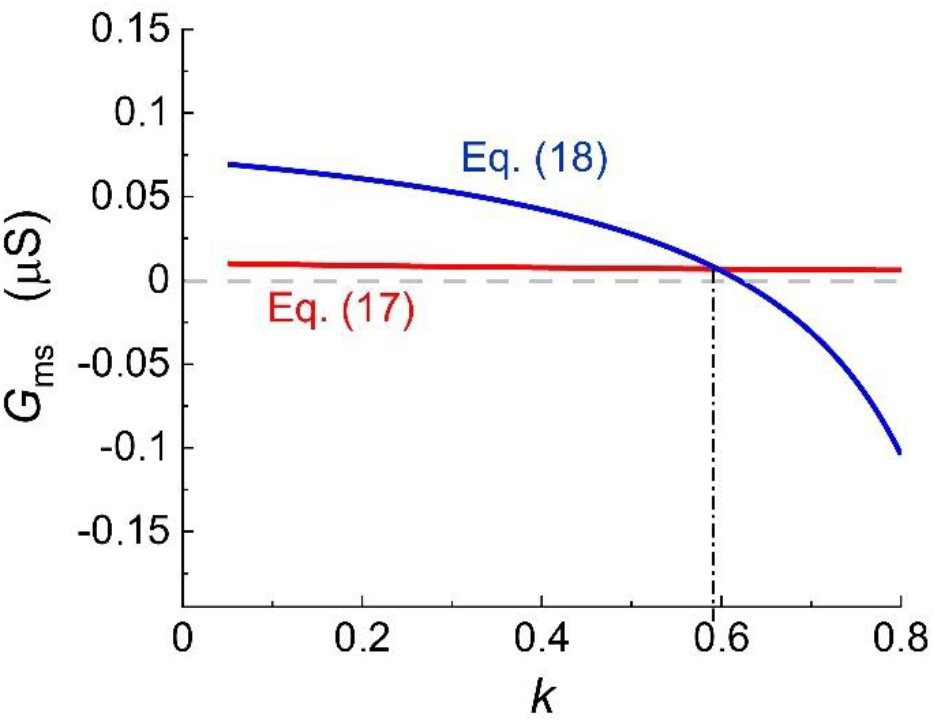
Graphic solution of the system of equations (17) and (18). The surface membrane conductivity *G*_ms_ was calculated as a function of *k* from Eq (17) (red) and Eq (18) (blue) in Matlab. The real values of *k* and *G*_ms_ were read at the point of intersection of the two curves.

The main objective of the proposed method was the evaluation of the membrane capacitances *C*_t_ and *C*_s_ and the fraction of tubular capacitance *f*_t_ = *C*_t_/(*C*_t_ + *C*_s_) as an estimate of the fraction of tubular membrane area *S*_t_/(*S*_t_ + *S*_s_). Hence, it was necessary to prove that these quantities calculated applying the proposed approach were independent on the values of other elements of electrical equivalent circuit (Fig 5). In this case, the coefficient *ɣ* was fixed at 0.7, while the resistance of the tubular membrane *R*_mt_ was variable to satisfy Eq (15). This required setting *R*_mt_ = *R*_ms_/(*ɣC*_t_/*C*_s_). First, we investigated the effect of changes in the access resistance *R*_a_ while maintaining the values of other parameters (except for the variable *R*_mt_) according to Table 2. The pre-set values of *C*_s_, *C*_t_, and *f*_t_ were well reproduced (Fig 5A, left) despite marked variations in the time courses of capacitive current (right). The correctness of the derived formulas was confirmed also at variable tubular membrane capacitance *C*_t_ settings (Fig 5B, left). The right panel illustrates the assessment of the coefficient *k* (in the way shown in Fig 4). The quantities *k* and *G*_ms_ varied with variable *C*_t_ (while *C*_s_ remained constant). Similarly, the capacitance values were well reproduced when the resistances *R*_ms_ and *R*_t_ were altered (Fig 5C). As mentioned above, in all these calculations the condition determining the interdependence of membrane resistances *R*_ms_ and *R*_mt_ (Eq (15)) was presumed to be met (illustrated for *γ* = 0.7). The error depending on the accuracy of the fitting process did not exceed 1%.

**Fig 5.**
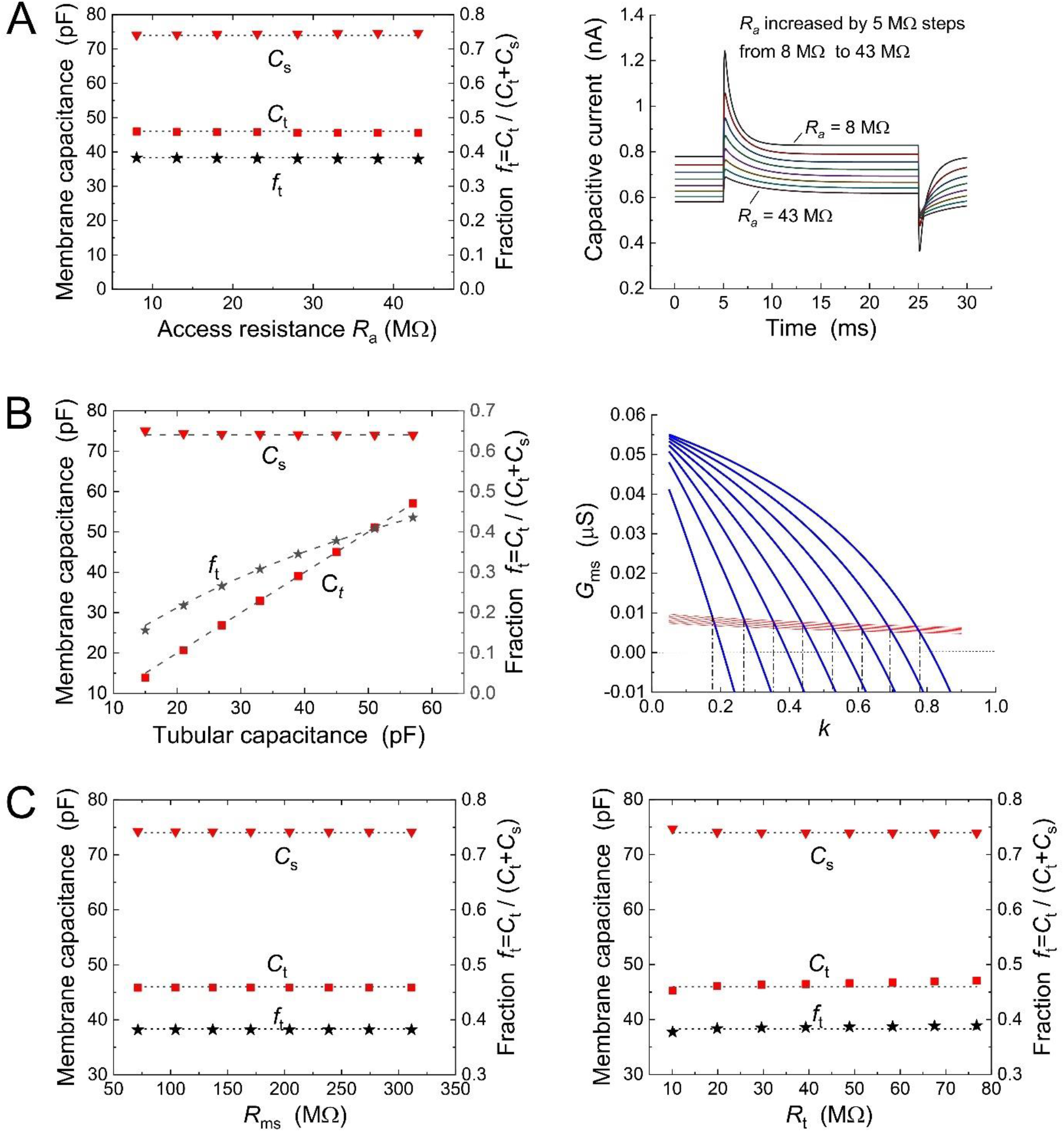
Verification of the proposed approach to determine the membrane capacitances *C*_s_, *C*_t_, and the fraction of tubular capacitance/area *f*_t_. Insensitivity of the calculated quantities: (A) to variations in access resistance *R*a (right panel shows time courses of the capacitive current), (B) to variations in tubular capacitance *C*t (right panel illustrates the determination of *k* = *C*_t_/*C*_s_ from Eq (17) and (18)), (C) to variations in surface membrane resistance *R*_ms_ (left) and resistance of the lumen of t-tubules *R*_t_ (right). Filled symbols: calculated values of *C*_s_, *C*_t_ = *k C*_s_ and *f*_t_ =*k*/(1+*k*); dotted lines: preselected values of *C*_s_ = 74 pF and *C*_t_ = 46 pF in A and C or preselected variation of *C*_t_ and thus *f*_t_ in B.

The choice of numerical values of all elements of the electrical equivalent scheme unambiguously determines the value of *γ* according to Eq (15) (pre-set *γ*-value). In real experiments on cardiac cells, the value of *γ* is unknown in advance and an estimate of *γ* (expected γ-value) is necessary to quantify the elements of the equivalent scheme. It is thus important to estimate the error caused by the difference between the expected and the pre-set *γ*-values. In Fig 6, the calculated tubular capacitance *C*_t_ and the fraction *f*_t_ are plotted as a function of the expected *γ*-value in the range 0.4 and 1.2. The calculated *C*_s_ does not depend on the *γ*-value and is not subject to error. If the values of all parameters were set according to Table 2 (except for *R*_mt_ ~ 345 MΩ corresponding to the pre-set *γ*-value 0.7), the error in the evaluation of *C*_t_ and *f*_t_ did not exceed 3% in the whole range of expected *γ*-values (Fig 6A). For comparison, Fig 6B shows that the error became negligible when the resistance of the tubular membrane was set to *R*_mt_ = *R*_ms_/(*γ C*_t_ /*C*_s_), so that the condition of Eq (15) was permanently satisfied.

**Fig 6.**
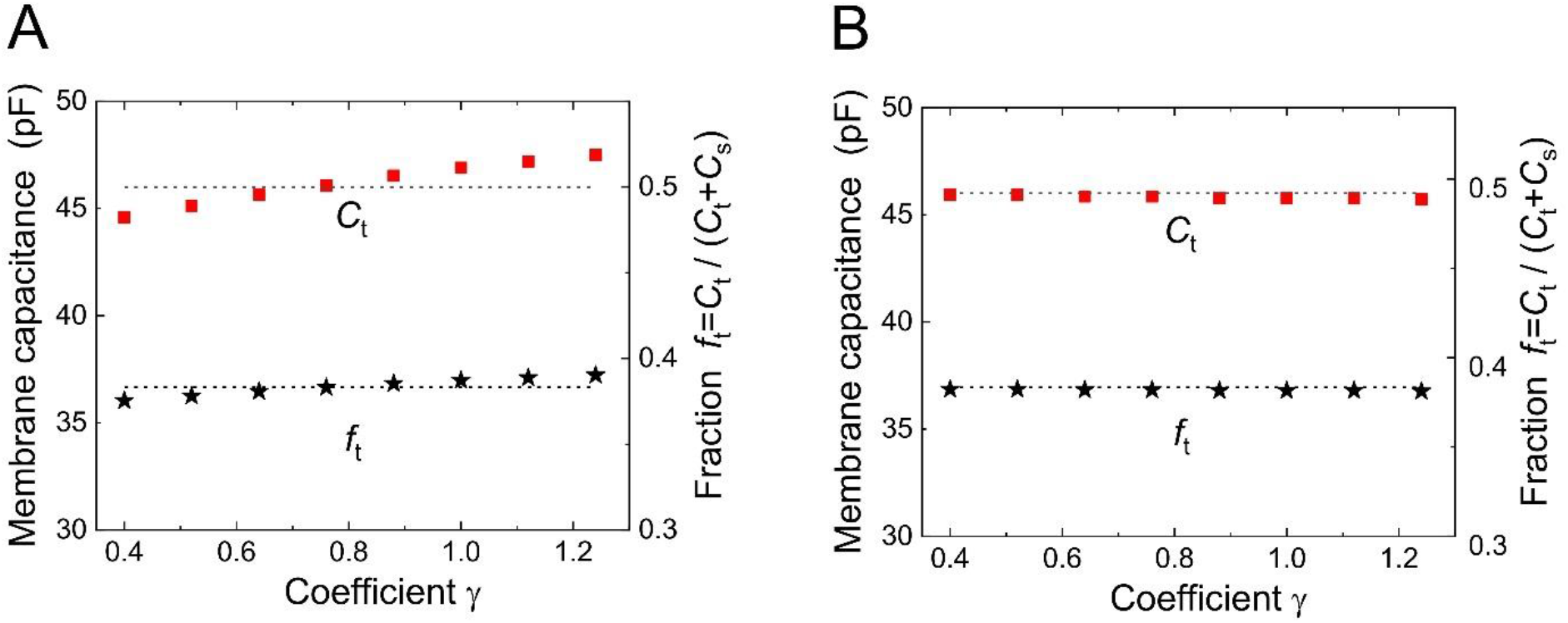
Estimation of the error in *C*_t_ and *f*_t_ determination caused by a difference between the expected *γ*-value and the real pre-set *γ* value (satisfying condition of Eq (15)). The expected *γ*-values ranged between 0.4 and 1.25. The calculated *C*_s_ (not shown) did not depend on the *γ*-value and was not subject to error. (A) Values of all parameters were set according to Table 2 (except for *R*_mt_=345 MΩ adjusting the pre-set *γ*-value to 0.7). The error in the evaluation of *C*_t_ and *f*_t_ did not exceed 3% over the entire range of expected *γ*-values. Note the zero error if the expected *γ* = 0.7. (B) For comparison, the resistance of the tubular membrane was set to *R*_mt_ = *R*_ms_/(*γ C*_t_/*C*_s_) so that the condition of Eq (15) has always been met. Dotted lines - preselected values of *C*_t_ and *f*_t_.

### Use of the method in experiments on ventricular cardiomyocytes

To examine changes of the membrane current caused by the isotonic sucrose solution, 2-s ramp-like membrane voltage from −160 to −40 mV and back was imposed at 0.1 Hz to the measured cell (Fig 7, upper panel). When Tyrode solution was replaced with sucrose solution, the inward current at a holding voltage of −80 mV was reversed (Fig 7, lower panel). The reversal membrane voltage, which was approximately −75 mV in Tyrode solution, was shifted to around −140 mV in sucrose solution. The recorded current probably corresponded to the potassium current *I*_K1_ as discussed later.

**Fig 7.**
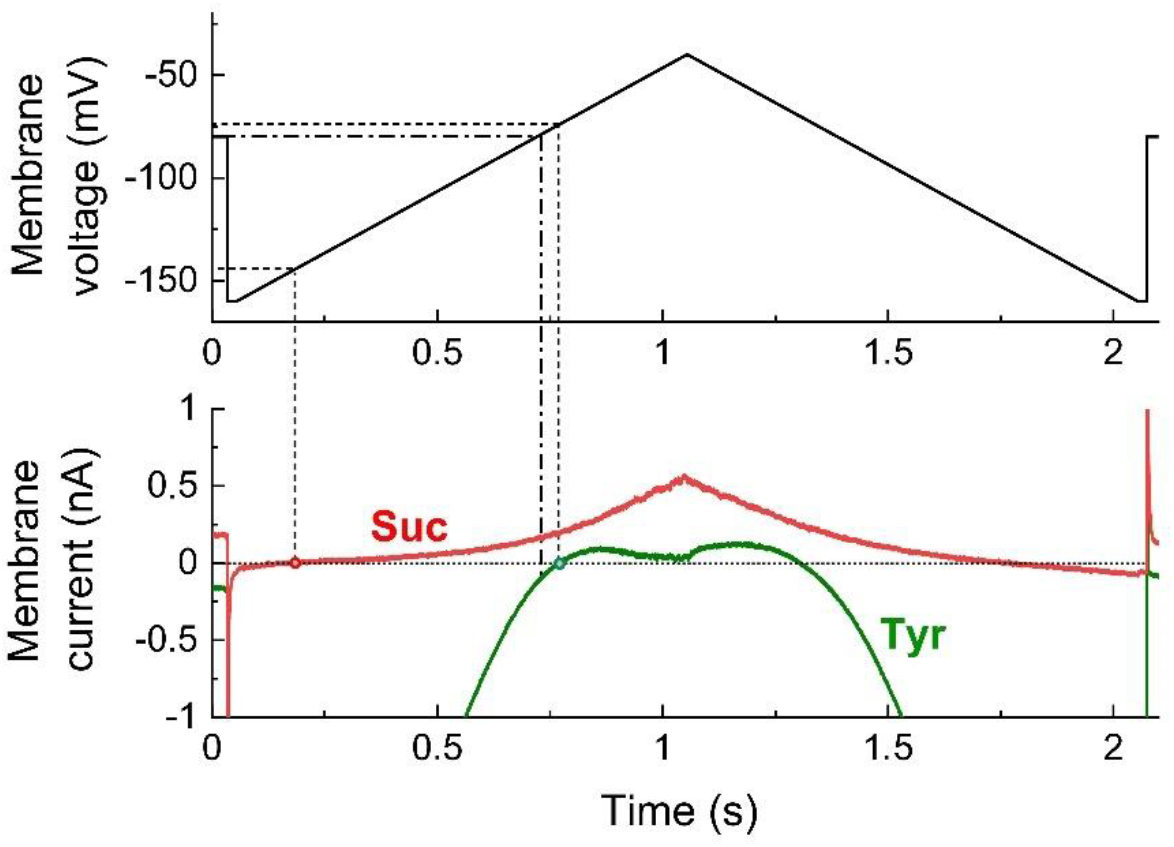
Comparison of steady-state current-voltage relationship in isotonic sucrose (Suc) and Tyrode (Tyr) solution. Membrane currents (lower panel) were recorded in response to changes in membrane voltage composed of 1 s ascending and 1 s descending ramp function in the range between −160 and −40 mV (upper panel). Note that the resting membrane voltage in Tyrode solution (−74 mV) was shifted to approximately −140 mV in sucrose solution (dotted lines). When Tyrode solution was replaced with sucrose solution, the inward current at the holding voltage of −80 mV reversed into an outward current (dash-dotted line).

The newly developed method was then tested in a pilot set of experiments on enzymatically isolated rat ventricular myocytes (*n* = 10). A train of 300 rectangular voltage steps (20 ms, 5 or 10 mV from the holding voltage of – 80 mV) was applied at 25 Hz to reach the steady-state. The last 50 current responses were averaged and evaluated. This procedure was repeatedly applied in the sucrose and Tyrode solution.

It was important to find out to what extent the resulting values of the main parameters depended on the estimation of the coefficient *γ*. The capacitance of the surface membrane *C*_s_= 72.27 ± 16.36 pF as calculated from Eq (10 and 11) did not depend on *γ*. The calculated values of capacitance and fraction of the tubular membrane increased slightly with *γ* growth. When setting *γ* = 0.7 and *γ* = 1.2, the values *C*_t_ = 40.6 ± 13.9 pF, *f*_t_ = 0.36 ± 0.08 and *C*_t_ = 43.7 ± 15.5 pF, *f*_t_ = 0.37 ± 0.08, respectively, were calculated.

Repetitive estimation of these parameters in the same cells was possible when the sucrose solution was washed out and applied again. Three separate applications of the sucrose solution resulted in similar values of *C*_m_, *C*_s_, and *C*_t_ as shown in a representative cell in Fig 8. As apparent, *C*_m_ was constantly below *C*_Tyr_ in the same cell (by ~18% on average; see Discussion for more details).

**Fig 8.**
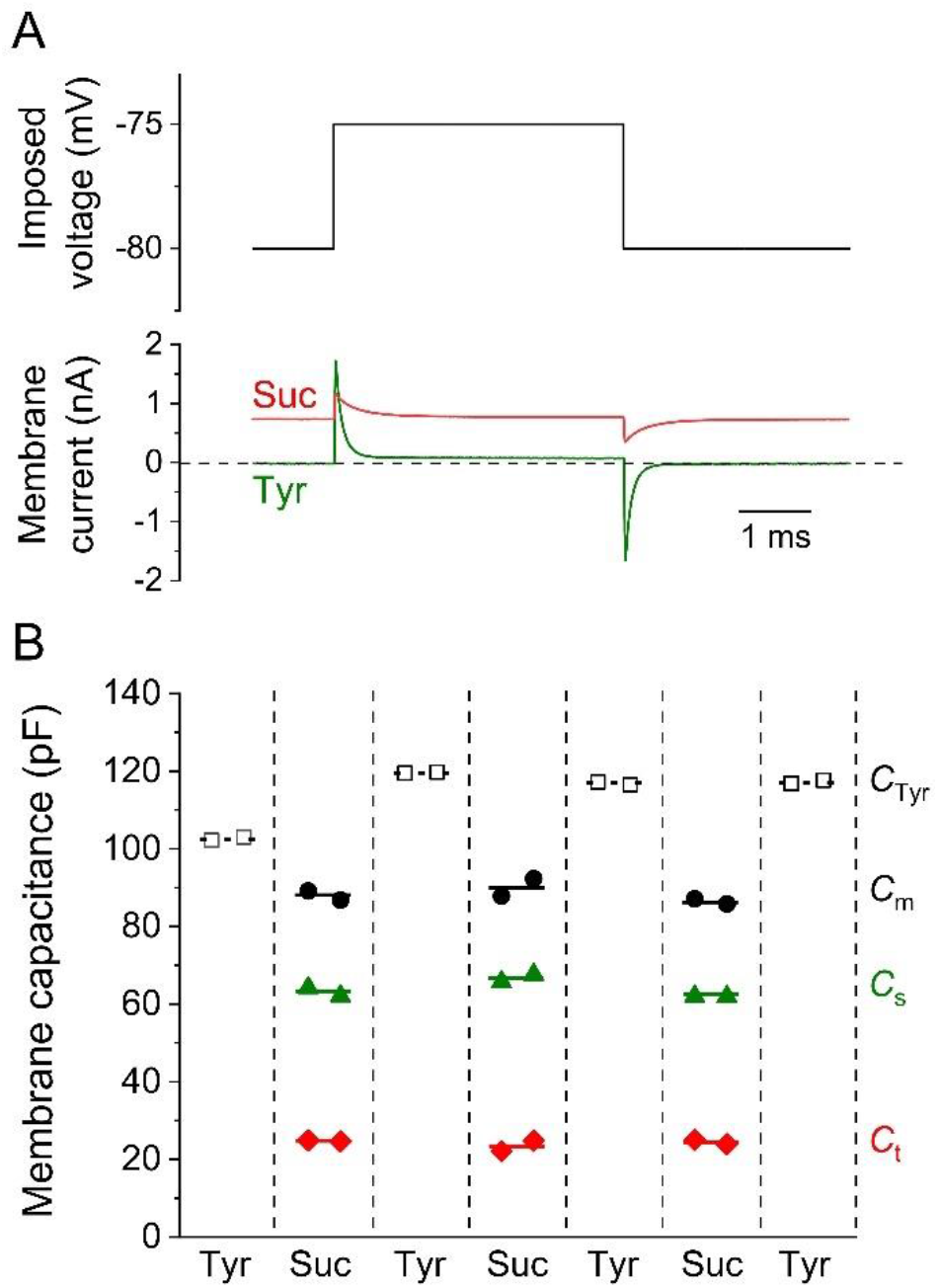
Experimental analysis of tubular and surface membrane capacitances using the newly developed method. (A) Scheme of the experimental protocol (upper panel) and representative recordings of the membrane current in Tyrode (Tyr) and sucrose (Suc) solutions (lower panel). (B) Tubular and surface membrane capacitance (*C*_t_, *C*_s_) during repetitive measurements compared with capacitance estimated in Tyrode solution (*C*_Tyr_) in a representative experiment. The cell was alternatively exposed to isotonic sucrose and Tyrode solution. Two subsequent evaluations were performed at the end of each steady-state application (50 consecutive current traces were averaged and fitted). Note that the total capacitance (*C*_m_ = *C*_s_ + *C*_t_) measured in the sucrose solution was reduced compared to *C*_Tyr_.

## Discussion

The present work aimed to formulate quantitatively and verify the idea that a substantial reduction in the electrical conductivity of the extracellular solution and the associated increase in lumen resistance of the t-system will allow us to quantify separately surface and tubular membrane capacitances (*C*_s_ and *C*_t_) from the results of double-exponential approximation of the capacitive current in cardiomyocytes. This approach was intended as an alternative to the known detubulation techniques with the aim to keep the cardiac cells intact and thus allow repeated measurements on the same cell. To suppress conductivity of the extracellular solution, we used the isotonic solution of sucrose.

Membrane current responses to small voltage-clamped rectangular impulses were evaluated to determine the elements of the model with lumped parameters (Fig 1B). Membrane capacitances (*C*_s_ and *C*_t_) are considered as indicators of membrane areas. Their separate determination is important because the tubular system membrane is functionally significantly different from the surface membrane (reviewed by Brette and Orchard [2]). The presence of two capacitances in combination with resistors implies bi-exponential current responses to the imposed steps of membrane voltage. However, in the case of cardiomyocytes in physiological solution, a low tubular lumen resistance causes that the magnitude of one of the two components is tiny and the corresponding time constant is so short that this component is indistinguishable. In contrast, both components could be distinguished in skeletal muscle fibers, which have a smaller diameter and therefore higher luminal resistance of the t-tubules [11].

In the proposed approach, the resistance of the lumen of the tubules as well as the resistance of the tubular membrane were temporarily substantially increased by the effect of an isotonic sucrose solution. A question arises as to the nature of the ionic current which remains after the replacement of the Tyrode’s solution with the solution containing only a residual amount of ions. To get a basic idea, we recorded the steady-state current-voltage relations using slow ramp pulses in isotonic sucrose and Tyrode solution for comparison (Fig 7). The reversal (zero current) voltage was shifted from around −70 mV in Tyrode to ~ −140 mV in sucrose solution. We considered *I*_K1_ to be likely responsible for residual ionic current in sucrose solution. The outward potassium current may then create a relatively stable thin peripheral layer adjacent to the membrane of enhanced K^+^ concentration which determines negative equilibrium voltage according to the Nernst equation. Considering the reversal voltage around −140 mV and K^+^ concentration in the pipette (~150 mM), the extracellular K^+^ concentration in this thin superficial layer could be roughly estimated to 0.5 mM during the sucrose application.

The main advantage of the proposed approach is the reversibility of the condition after exposition to a low conductance solution. The measurements in sucrose solution may be several times replaced by measurement in physiological (Tyrode) solution as illustrated in Fig 8. In comparison with the irreversible measurement using detubulation techniques, the proposed approach allows repetitive measurements in the same cell and application of the paired tests. The method could also be useful for separate monitoring of short-term changes in *C*_t_ and *C*_s_ caused e.g. by osmotic shocks [7, 12].

The lumped-element model used to describe the membrane system is simplistic as well as the more complex distributed model. The arrangement of the network of interconnected tubules imaged by microscopic methods in cardiac cells [13–15] differs from parallel arranged transverse tubules described by cable equations. Moreover, the use of the lumped model is supported by experiments indicating that in rat cardiomyocytes, the tubular length constant λ = (*r*_mt_ / *r*_t_)^0.5^ is one order of magnitude larger than the cellular radius [16]. The symbols *r*_mt_ and *r*_t_ denote tubular membrane resistance [Ω m] and resistance of the lumen [Ω m^−1^] per unit of radial tubular length, respectively. This suggests that the drop of membrane voltage along the radial tubules can be regarded negligible so that the tubules are virtually uniformly polarized.

Another simplification is the replacement of voltage-dependent membrane resistances by constants, which corresponds to a linear approximation of the current-voltage relation in the vicinity of the holding voltage (constant slope conductance). Nevertheless, this limitation is minimized by selecting a sufficiently small voltage step for capacitance measurement.

The cell membrane capacitance has been reported to be reduced in skeletal muscle fibers exposed to solutions of low ionic strength [17]. Our preliminary results showed an average decrease of the total membrane capacitance in isotonic sucrose solution expressed by the sum *C*_m_ = *C*_s_ + *C*_t_ compared to the capacitance *C*_Tyr_ measured in the Tyrode solution by ~18%. Yet if the decrease in *C*_s_ and *C*_t_ were the same, the coefficient of the fraction of tubular capacitance *f*_t_ (as an indicator of membrane areas ratio) would remain unchanged. Moreover, *C*_m_ and *C*_Tyr_ values are available from repeated measurements on a given cell. Thus, the *C*_s_ and *C*_t_ values can be easily corrected for decreases caused by sucrose solution.

To explain different estimates of t-tubule membrane fraction obtained using the detubulation methods (~32%) and optical measurements (up to 65%), Pásek et al. (2008b) suggested that the specific capacitance of the t-tubule membrane may be lower than that of the surface membrane, possibly because of high cholesterol content. If quantified, this effect can be taken into account in Eq (2).

The introduction of the coefficient *γ* (i.e., the proportionality constant between the ratio of membrane capacitances and membrane conductivities) plays an important role in the presented theoretical approach. According to Fig 6, the error due to the deviation between the expected and the actual value of *γ* was less than 3% and the difference in *f*_t_ estimated from measurements on rat cardiomyocytes at the expected *γ*-values of 0.7 and 1.2 was approximately 4%. Another way to solve the problem of determining the capacitance *C*_t_ according to Eq (14) without estimating the ratio *γ* is the substitution of parallel combination *R*_mt_ || *R*_t_ for *R*_ms_ || *R*_t_, which is justified if *R*_mt_ >> *R*_t_ and *R*_ms_ >> *R*_t_. This procedure will be a subject of another study supplemented by applicability criteria and detailed experimental results.

## Material and methods

### Experiments on rat ventricular cardiomyocytes

Cardiomyocytes were enzymatically isolated as described before (Bébarová et al. 2005) from right ventricles of adult male Wistar rats (275 ± 50 g) anaesthetized by intramuscular administration of a mixture of tiletamine and zolazepam (65 mg kg^−1^; Zoletil® 100 inj.), and xylazine (20 mg kg^−1^; Xylapan® inj.). The experiments were carried out with respect to recommendations of the European Community Guide for the Care and Use of Laboratory Animals; the experimental protocol was approved by the Local Committee for Animal Treatment at Masaryk University, Faculty of Medicine, and by the Ministry of Education, Youth and Sports (permission No. MSMT-29203/2012-30 and MSMT-33846/2017-3).

Tyrode solution with the following composition was used both during the dissociation procedure and to perfuse myocytes during the measurements (in mM): NaCl 135, KCl 5.4, MgCl_2_ 0.9, HEPES 10, NaH_2_PO_4_ 0.33, CaCl_2_ 0.9, glucose 10 (pH was adjusted to 7.4 with NaOH). CoCl_2_ (2 mM; 1 M stock solution in the deionized water) was used for inhibition of *ICa*. The patch electrode filling solution contained (in mM): L-aspartic acid 130, KCl 25, MgCl_2_ 1, K_2_ATP 5, EGTA 1, HEPES 5, GTP 0.1, Na_2_-phosphocreatine 3 (pH 7.25 adjusted with KOH). Sucrose solution (3.2 M) was prepared by adding the deionized water into the sucrose (purity ≥ 99.5%).

Single rod-shaped cardiomyocytes with well-visible striations were used for the recordings applying the whole-cell patch-clamp technique in the voltage-clamp mode at room temperature (23 ± 1°C) using the Axopatch 200B equipment and pCLAMP 10.2 software. The patch pipettes were pulled from borosilicate glass capillary tubes and heat polished on a programmable horizontal puller; the resistance of the filled glass electrodes was below 1.5 MΩ to keep the access resistance as low as possible. For experimental protocols, please see the Results; the membrane currents (recorded using the ramp and rectangular protocol) were digitally sampled at 100 and 200 kHz, respectively. The recorded data were evaluated as described in the Results using the following software: Clampfit (v.10.2), MATLAB (v.R2017a), and Origin (v.2015).

The accuracy of tubular capacitance determination depends on how thoroughly the tubular system is washed with sucrose solution. The jet pipes for rapid exchange of solutions must be reliably directed at the cell under examination. The magnitude of the change in access resistance can be used as a criterion. A part of the tubular system may be less accessible to sucrose solution if the cell lies at the bottom of the chamber. It is best to lift the cell, which may be however risky. Incomplete solution exchange will affect the ratio of magnitudes and time constants of both components of the analyzed part of the capacitive current. The unacceptably low resistance of the t-system lumens will also affect the ratio *R*_1_/*R*_2_ of the resistances calculated according to Eq (12) and (13). To decide whether a given measurement is acceptable and can be included in the overall evaluation, we set the following criteria:

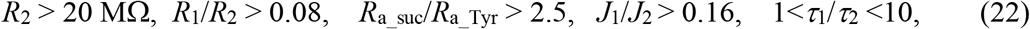

where *R*_a_suc_, and *R*_a_Tyr_ are access resistances in sucrose and Tyrode solution, *τ*_1_ refers to the longer of the two time constants.

In all experiments, the capacitive current was approximated by a bi-exponential function using the Matlab Clampfit software. The resulting values of the parameters *J*_1_, *J*_2_, *J*_∞,2_, *J∞*,_1_, *τ*_1_, and *τ*_2_ were then transferred to the software S2_evaluation.mlx provided with all computational relationships and necessary procedures for quantification of parameters of the electrical equivalent scheme. The results of measurements that met the criteria (22) are included in the S2_evaluation.mlx executable file available in the Supporting Information.

## Supporting information

Supplement S1_verification

Supplement S2_evaluation

## Supporting information

The following files written in Matlab language were transferred to the Matlab Live Editor for clarity.

The file “**S1_verification.mlx**” allows to perform verification ‘computer experiments’ with optional values of the elements of the electrical equivalent circuit as described in the section of Results “Model verification of the theory”. This software allows to perform verification of the theoretical background with optional values of the elements of the electrical equivalent circuit (for details see section of Results “Model verification of the theory”). The ‘computational experiments’ were designed to mimic the real experiments in isolated cells.

The user has the following options: (i) If F=0, the coefficient *γ is* determined by the selected values of the equivalent circuit parameters to meet the condition of Eq (15). If F=1, *γ* is to be selected in the interval between 0.4 and 1.2. (ii) The values of the parameters of electrical equivalent circuit are preset according to Table 2 but can be changed by the user in a limited range. (iii) After running the file, graphs of the solution corresponding to Figs. 2–4 will appear on the right in the Live Editor window with tables comparing the preset and calculated parameter values.

The file “**S2_evaluation.mlx**” includes (i) the data from all experiments evaluated in this study as preliminary results and (ii) calculations based on the derived mathematical formulas for quantification of electrical equivalent scheme parameters resulting from the data of the selected experiment. In each measurement, the vector MM contains the numeric values of the capacitive current fit in the following order: the magnitude and the time constant of the first exponential component (A1 in nA and T1 in ms), the magnitude and the time constant of the second exponential component (A2 in nA and T2 in ms), and the steady-state values of the current at the holding and the imposed impulse voltage (JO1 and JO1 in nA). After conversion to the notation given in the Results section these are quantities *J*_1_, *τ*_1_, *J*_2_, *τ*_2_, *J*_∞,1_, and *J*_∞,2_. However, since the A1 and A2 are determined at the time delayed after the beginning of the voltage step by the time Dt (ms), *J*_1_ and *J*_2_ are calculated in the following program steps by extrapolation of A1 and A2 to the beginning of the voltage step. To calculate the elements of the electric equivalent circuit, it is necessary to run the program after uncommenting the selected experimental result. The right column of the Live Editor window shows the graphical solution of the Eq (17) and (18) and tables of the resulting numerical values of the elements of the equivalent circuit.

Both described files have been converted to pdf format and are available as supplements S1_verification.pdf and S2_evaluation.pdf.

## Abbreviations

*C*_m_: Total membrane capacitance
*C*_t_: Tubular membrane capacitance
*C*_s_: Surface membrane capacitance
*d*: Membrane thickness
*ε*: Membrane permittivity
*γ*: Coefficient of proportionality between ratio of membrane capacitances and membrane conductivities
*J*: Membrane current
*J*_1_, *J*_1_: Magnitudes of exponential components of the capacitive current
*J*_∞,1_, *J*_∞,2_: Steady-state currents at the membrane voltages *U*_1_ and *U*_2_
*k*: *C*_t_/*C*_s_ ratios
*R*_a_: Access resistance
*R*_ms_, *R*_mt_: Membrane resistances
*R*_t_: Tubular system lumen resistance
*S*_s_, *S*_t_: Surface and tubular membrane area
*τ*_1_, *τ*_2_: Time constants of exponential components of the capacitive current
*U*_1_, *U*_2_: Imposed levels of membrane voltage
*U*_s_, *U*_t_: Surface and tubular membrane voltage

## Author contributions

Conceptualization: JŠ.

Formal analysis: JŠ, MŠ, and MB.

Investigation: OŠ and MB.

Methodology: JŠ.

Software: JŠ.

Writing – original draft: JŠ, MŠ, and MB.

Writing – review and editing: JŠ, MŠ, and MB.

