## Supplement S1_verification for "A new approach to the determination of tubular membrane capacitance: passive membrane electrical properties under reduced electrical conductivity of the extracellular solution"

### CALCULATION OF PARAMETERS OF ELECTRICAL EQUIVALENT CIRCUIT

J. Šimurda, M. Šimurdová, O. Švecová, M. Bébarová

#### Model verification of the theoretical background

This  software allows to perform verification of the theoretical background with optional values of the elements of the electrical equivalent circuit (for details see section of Results “Model verification of the theory" in the main text). The 'computational experiments' were designed to mimic the real experiments in isolated cells.

The user has the following  options:

- If F=0, the coeffitient $\gamma$ *is*  determined by the selected values of the equivalent circuit parameters to meet the condition of Eq (15). If F=1, $\gamma$  is to be selected in the interval I between 0.4 and 1.2.
- The values of the parameters of electrical equivalent circuit are preset according to Table 2 in the main text but can be changed by the user (in a limited range).
- After running the file, graphs of the solution corresponding to figures 2 - 4 in the main text will be displayed with tables comparing the preset and calculated parameter values.

```
%close all
%clear all
global Ums Umt U1 U2 Too T To tau
global Tt Ts kt kst ket kUt kUs
```

Option:  if F1=0,  gamma is set to meet condition of Eq. 15;  if F=1, gamma can be set arbitrarily in the interval between 0.4 and 1.2

```
Fl=0;
% Fl=1; gamma=0.7;
```

Selected values of the elements of electrical equivalent circuit:

```
% basic choice (nF; MO; mV)
Cs=0.074;     Ra=12.5;
Ct=0.046;     Rt=15;
Ums=-160;     Rms=150;
Umt=-160;     Rmt=241;


% altered values
% Cs=;
% Ct=0.015;         % smaller Ct must be combined with high Rt
% Ra=38;
```

```
% Rt=;
% Rms=;
% Rmt=360;
% Ums=;
% Umt=;
```

Calculation of values of the derived parameters

```
Tt=Rt*Ct; Ts=Ra*Cs; kt=1+Rt/Rmt; kst=1+Ra/Rms+Ra/Rt; ket=Ra/Rt;
kUt=Umt*Rt/Rmt; kUs=Ums*Ra/Rms; ft=Ct/(Cs+Ct);
```

Gamma coefficient

```
if Fl==0
ga=(Cs/Ct)*(Rms/Rmt);
else
    ga=gamma;
end
```

Time constant related to the rising and falling edge of the imposed rectangular pulse

```
tau=0.05;
```

The fitting starts in the instant delayed by *D*t (ms) after the onset of the depolarization step.

```
Dt=0.4;
```

Stimulation protocol

```
T=20; To=30; Too=5;
% T-duration of the voltage impulse, To-time period, % Too-delay

U1=-80; U2=-75 ;
% holding and impulse voltage
```

SOLUTION OF DIFFERENTIAL EQUATIONS

Initial conditions

```
Us0=(kt*kUs+ket*kUt+kt*U1)/(kst*kt-ket);
Ut0=(kUs+kst*kUt+U1)/(kst*kt-ket);
u0=U1;
```

Computation parameters

```
Tstart=0; Tend=30; InitialStep=0.0000001; MaxStep=0.005; TOL=1e-4;
```

Solution

```
options=odeset('InitialStep',InitialStep,'MaxStep',MaxStep);
[t,y]=ode15s(@model,[Tstart Tend],[Us0; Ut0; u0],options);
```

```
Us=y(:,1);
Ut=y(:,2);
u=y(:,3);

J=(u-Us)/Ra;

plot(t, u)
title('Imposed voltage')
xlabel('Time (ms)')
ylabel('Membrane voltage (mV)')
axis([0 30 -81 -74])
grid on
```

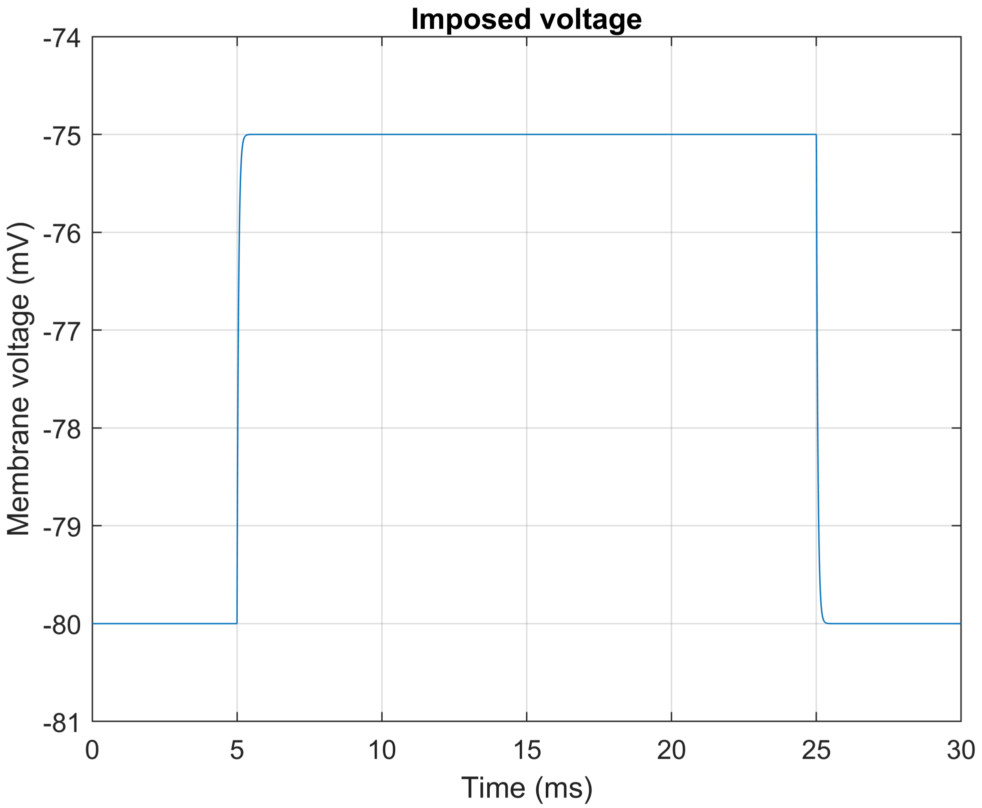


```
plot(t, Us, t, Ut)
title('Membrane voltage')
xlabel('Time (ms)')
ylabel('Membrane voltage (mV)')
legend({'surface','tubular'})
%  axis([0 30 -94 -76])
grid on
```

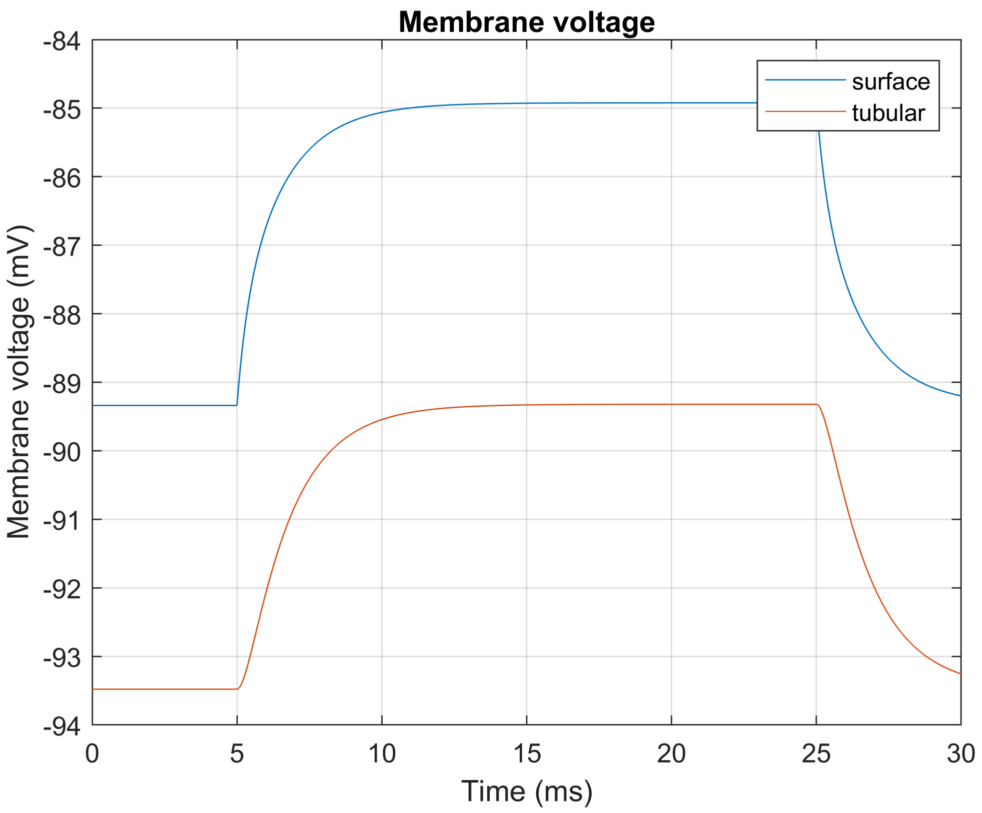


```
plot(t,J,'-')
axis([0 30 0.4 1.15])
title('Capacitive current')
xlabel('Time (ms)')
ylabel('Current (uA)')
grid on
```

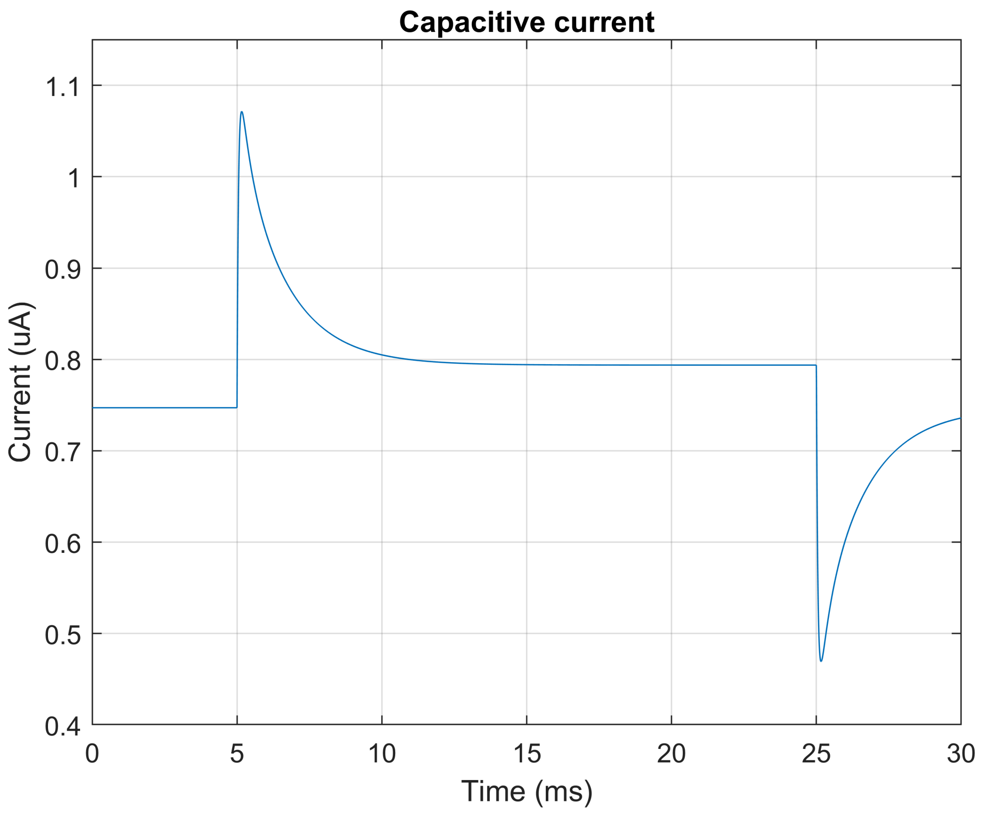


Separation of part of capacitive current for fitting by a bi-exponential function

```
fi=find((abs(t-Too-Dt))<1e-2);
fe=find((abs(t-24.9))<1e-2);
gi=fi(length(fi));
ge=fe(length(fe));
g=gi:ge;
```

Separated part of the capacitive current

```
tt=t(g);
II=J(g);
```

Fitting procedure using Matlab fitting tool

```
 f=fit(tt,II,'J1*exp(-(x-5)/T1)+J2*exp(-(x-5)/T2)+J02', 'StartPoint',...
     [0.251 1.4 0.08 0.34 1.1],...
     'Lower',[0 0 0 0 -Inf]);
```

Parameters resulting from the fitting procedure

```
J1=f.J1;
J2=f.J2;
T1=f.T1;
T2=f.T2 ;
J02=f.J02;
```

```
J01=J(1);
```

Separated part of the capacitive current overlaid by its bi-exponential fit

```
fit_J=J1*exp(-(tt-5)/T1)+J2*exp(-(tt-5)/T2)+J02;

plot(tt, II, tt, fit_J)
% axis([5 25 0.7 1.3])
title('Capacitive current overlaid by its bi-exponential fit')
xlabel('Time (ms)')
ylabel('Current (uA)')
grid on
```

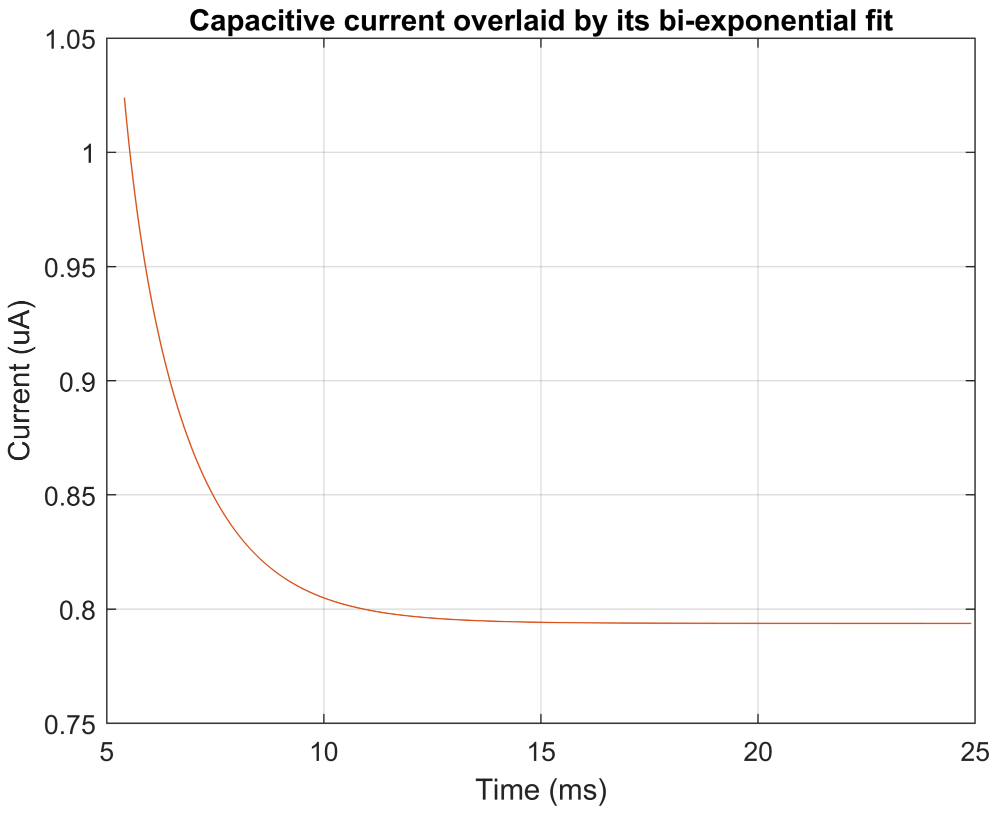


Calculation of some parameters listed in Table 1

The symbol f indicates that the parameter is derived from a bi-exponential fit of capacitive current

```
Raf=(U2-U1)/(J1+J2+J02-J01);
Tsf=(J1+J2+J02-J01)*T1*T2/(T1*J2+T2*J1);
Csf=Tsf/Raf;
a=(J1+J2)/(J1+J2+J02-J01);
b=(T1^2*J2+T2^2*J1)*Tsf/((T1*J2+T2*J1)*T1*T2);
R1=Ra/(b-1);
R2=Ra*a/(1-a);
Re12=Ra*(T1*J2+T2*J1)/((J1+J2)*Tsf);
```

Calculation of the parameter $k=Ct/Cs$:

*Gms* = 1/*R*ms expressed from Eq.17 as a function of the variable *k* (denoted *kk*) is denoted as *Gms*1

```
kk=0.001:0.001:1.8;
Rms1=(R1+R2+ga*kk*(R2-R1)+((R1+R2+ga*kk*(R2-R1)).^2-4*R1*R2).^0.5)/2;
Gms1=1./Rms1;
```

*Gms* = 1/*R*ms expressed from Eq.18 as a function of the variable *k* (denoted *kk*) is denoted as *Gms*2

```
Rms2=((ga*kk-1)*R1*Re12)./(kk*R1-Re12);
Gms2=1./Rms2;
```

Graphic solution of the system of Eqs 17 and 18

```
plot(kk,Gms1,kk,Gms2)
axis([0 0.9 -0.1 0.1])
title('Graphic solution')
xlabel('kk')
ylabel('Gms (uS)')
legend({'Gms1','Gms2'})
grid on
```

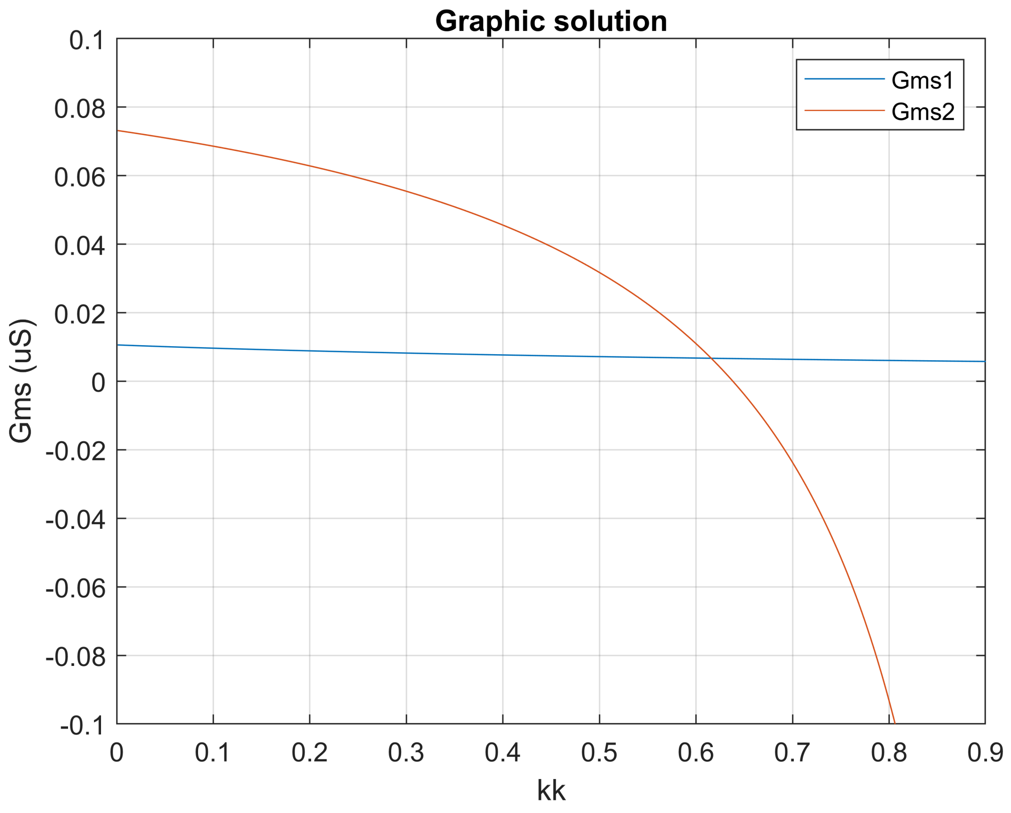


```
% hold on
```

Solution of the system of Eqs.17 and 18 to calculate *k*

```
q=find(abs((Gms2-Gms1))-min((abs(Gms2-Gms1)))==0);
if q>1
Gmsf=Gms2(q(1)); Rmsf=1/Gmsf;
k=kk(q);
```

Calculation of the remaining parameters included in Table 1

```
Ctf=k*Csf;
ftf=Ctf./(Csf+Ctf);
Cmf=Csf+Ctf;
Rtf=R1*Rmsf/(Rmsf-R1);
Rmtf= Rmsf/(ga*k);
 else
     k=NaN; Csf=NaN; Raf=NaN; Rtf=NaN; Rmsf=NaN; Ctf=NaN; ftf=NaN; Rmtf=NaN;
    str=('out_of_range')
 end

 if q==length(kk)
 k=NaN; Csf=NaN; Raf=NaN; Rtf=NaN; Rmsf=NaN; Ctf=NaN; ftf=NaN; Rmtf=NaN;
    str=('out_of_range')
 end
```

Comparison between selected basic parameters and parameters determined according to the proposed method

```
Cs_f=1000*[Cs;Csf]; Ct_f=1000*[Ct;Ctf]; ft_f=[ft;ftf]; Ra_f=[Ra;Raf]; Rt_f=[Rt;Rtf];...
    Rms_f=[Rms;Rmsf]; Rmt_f=[Rmt;Rmtf];
k
```

```
k = 0.6160
```

```
ga
```

```
ga = 1.0013
```

```
Tab1=table(Cs_f, Ct_f, ft_f)
```

Tab1 = 2×3 table

| | Cs_f | Ct_f | ft_f |
|---|---|---|---|
| 1 | 74.0000 | 46.0000 | 0.3833 |
| 2 | 74.2097 | 45.7132 | 0.3812 |

```
Tab2=table(Ra_f, Rt_f)
```

Tab2 = 2×2 table

| | Ra_f | Rt_f |
|---|---|---|
| 1 | 12.5000 | 15.0000 |
| 2 | 12.5339 | 15.0203 |

```
Tab3=table(Rms_f, Rmt_f)
```

Tab3 = 2×2 table

| | Rms_f | Rmt_f |
|---|---|---|
| 1 | 150.0000 | 241.0000 |
| 2 | 150.7944 | 244.4873 |

```
%Ur=U2-J02*Raf/(1-a)%; Ur=U1-J01*Raf/(1-a)
```

Model - solution of differential equations

```
function dydt=model(t,y)
Us=y(1);
Ut=y(2);
u=y(3);

global  U1 U2 Too T To tau
global Tt Ts kt kst ket kUt kUs

if t<Too
   U=U1;
elseif mod(t-Too,To)<=T
    U=U2;
else
    U=U1;
end

dUs=(1/Ts)*(-kst*Us+ket*Ut+kUs+U);
dUt=(1/Tt)*(Us-kt*Ut+kUt);
du=(U-u)/tau;

dydt=[dUs; dUt; du];
end
```
