## Supplement S2_evaluation for "A new approach to the determination of tubular membrane capacitance: passive membrane electrical properties under reduced electrical conductivity of the extracellular solution"

### CALCULATION OF PARAMETERS OF ELECTRICAL EQUIVALENT CIRCUIT

J. Šimurda, M. Šimurdová, O. Švecová, M. Bébarová

The experiments on rat ventricular cardiomyocytes and their evaluation were performed using Axon Instruments equipment (Axopatch 200B) and associated software (Clampex and Clampfit). Membrane currents were recorded in response to the imposed depolarizing steps of membrane voltage. Evaluated are results of capacitive current fit by a sum of two exponential functions obtained by the least squares method in the Clampfit software.

*Experimental results from rat ventricular cardiomyocytes*

In each evaluated cell, the vector MM contains numeric values of the bi-exponential capacitive current fit in the following order:

Magnitude of the first exponential component *A*1 (nA);
Time constant of the first exponential component *T*1 (ms);
Magnitude of the second exponential component *A*2 (nA);
Time constant of the second exponential component *T*2 (ms);
Steady state value of current at holding voltage *J*O1 (nA);
Steady state value of current at the level of imposed step *J*O2 (nA).

The capacitive currents in response to 30 ms steps of the membrane voltage from *U*1= -80 mV to *U*2=-70 mV were fitted starting at the instant delayed after the beginning of the voltage step by the time *D*t (given in ms for each measured cell).

To run the file, it is necessary to uncomment the selected measurement.

```
% clc
% clear all

% Parameters from cell number 13205003 - komorové:
MM=[0.471649  1.55114 0.0975374  0.337172 0.4533  0.54379];
Dt=[0.2];

% % Parameters from cell number 13205004 - komorové:
% MM=[0.274036  2.13174  0.0520276  0.702072  0.3878  0.52239];
% Dt=[0.2];

% % Parameters from cell number 13205005 - komorové:
% MM=[0.30438  1.94682 0.0563664  0.499677  0.4291  0.567612];
% Dt=[0.2];

% % Parameters from cell number 132050010 - komorové:
% MM=[0.260789 1.88889 0.0663303  0.41934  0.784  0.978775];
% Dt=[0.2];
```

```
% % Parameters from cell number 132050011 - komorové:
% MM=[0.249585 1.91284 0.0633152  0.435593  0.8845  1.06334];
% Dt=[0.2];

% % Parameters from cell number 132050018 - komorové:
% MM=[0.154706 3.37461 0.199835  1.07159  0.9799  1.09261];
% Dt=[0.2];

% % Parameters from cell number 132050019 - komorové:
% MM=[0.271131 2.03963 0.0640055  0.416185  0.9009  1.03691];
% Dt=[0.2];

% % Parameters from cell number 132050044 - komorové:
% MM=[0.415116 1.4444 0.708754  0.487548  0.1773  0.237572];
% Dt=[0.2];

% % Parameters from cell number 132050052 - komorové:
% MM=[0.511943 1.72169  0.261467  0.414618  0.0602  0.109167];
% Dt=[0.1];

% %Parameters from cell number 132050054 - komorové:
% MM=[0.533545 1.53526 0.28271  0.390551  0.0402  0.0816154];
% Dt=[0.1];

% % Parameters from cell number 132050055 - komorové:
% MM=[0.495943 1.5336 0.314228  0.386779  0.0871  0.13702];
% Dt=[0.1];

% % Parameters from cell number132050063 - komorové:
% MM=[0.278442 2.75795  0.147132  0.784501  0.205  0.255075];
% Dt=[0.1];

% % Parameters from cell number 132050064 - komorové:
% MM=[0.37155  2.05961 0.104008  0.339378  0.1817  0.227513];
% Dt=[0.1];

% % Parameters from cell number 132050065 - komorové:
% MM=[0.330961  2.35039 0.148077  0.54016  0.1593  0.22128];
% Dt=[0.1];

% % Parameters from cell number 13219002 - komorové:
% MM=[0.159365 4.962  0.179969  1.21121  0.4827  0.532434];
% Dt=[0.1];

% % Parameters from cell number 13219003 - komorové:
% MM=[0.0899244  3.05211  0.0857916  0.878286  0.3452  0.406953];
% Dt=[0.2];

% % Parameters from cell number 13219010 - komorové:
% MM=[0.102693 2.526494  0.530023  0.603774  0.4342  0.552734];
% Dt=[0.2];
```

```
% % Parameters from cell number 13219011 - komorové:
% MM=[0.081057  3.490718  0.088733  1.015256  0.4774  0.544944];
% Dt=[0.2];

% % Parameters from cell number 13219021 - komorové:
% MM=[0.31474  1.99725 0.0376091  0.347416  0.8214  0.988587];
% Dt=[0.2];

% % Parameters from cell number 13219022 - komorové:
% MM=[0.291017 2.10102  0.0406958  0.335005  0.6925  0.856385];
% Dt=[0.2];

% % Parameters from cell number 13219026 - komorové:
% MM=[0.248671 2.15512 0.0255433  0.300621  0.817  1.00674];
% Dt=[0.2];

% % Parameters from cell number 13219033 - komorové:
% MM=[0.250223  2.13004 0.0351753  0.436589  0.272  0.315922];
% Dt=[0.2];

% % Parameters from cell number 13219039 - komorové:
% MM=[0.245475 2.09377  0.0336863  0.375736  0.2043  0.247449];
% Dt=[0.2];

% % Parameters from cell number 13219045 - komorové:
% MM=[0.501547 1.68712 0.524853  0.546366  0.1779  0.201546];
% Dt=[0.2];

% % Parameters from cell number 13219046 - komorové:
% MM=[0.269419 2.3685 0.0419532  0.368823  0.1886  0.254732];
% Dt=[0.2];

% % Parameters from cell number 13219055 - komorové:
% MM=[0.258325 2.22438 0.0325364  0.319549  0.2311  0.294777];
% Dt=[0.2];

% % Parameters from cell number 13219082 - komorové:
% MM=[0.38664  1.7553  0.0553337  0.3097  0.2626  0.32029];
% Dt=[0.2];

% % Parameters from cell number 13219085 - komorové:
% MM=[0.300362 2.00766  0.0531527  0.34862  0.1823  0.245612];
% Dt=[0.2];

% % Parameters from cell number 13219091 - komorové:
% MM=[0.328189 1.78957  0.0457228  0.288263  0.4011  0.482341];
% Dt=[0.2];

% % Parameters from cell number 13416005 - komorové:
% MM=[0.372380 2.072792  0.162790  0.474466  0.3065  0.342990];
% Dt=[0.3];

% % Parameters from cell number 13416006 - komorové:
```

```
% MM=[0.226340 3.318573  0.188345  0.675546  0.1118  0.133269];
% Dt=[0.3];

% % Parameters from cell number 13416019 - komorové:
% MM=[0.349075  1.554379  0.209296  0.574599  0.2024  0.228173];
% Dt=[0.22];

% % Parameters from cell number 13416021 - komorové:
% MM=[0.253079 2.133046  0.198951  0.618157  0.15  0.171172];
% Dt=[0.25];

% % Parameters from cell number 13416026 - komorové:
% MM=[0.223062 1.743620  0.362101  0.681484  0.3075  0.348136];
% Dt=[0.3];

% % Parameters from cell number 13416027 - komorové:
% MM=[0.157327 2.576585  0.257658  0.925235  0.1861  0.216387];
% Dt=[0.3];

% % Parameters from cell number 13416029 - komorové:
% MM=[0.13654  3.10727 0.264577  0.957714  0.1603  0.186126];
% Dt=[0.3];

% % Parameters from cell number 13416034 - komorové:
% MM=[0.273188 1.933865  0.091846  0.603048  0.2686  0.309869];
% Dt=[0.3];

% % Parameters from cell number 13416035 - komorové:
% MM=[0.297321 1.878515  0.100794  0.482217  0.1741  0.203537];
% Dt=[0.25];

% % Artificial cell (from suppl_1_verification.mlx)
% MM=[0.1794  0.3382  0.5357  1.6285 0.6687  0.7522];
% Dt=[0];
```

CALCULATIONS

Coefficient gamma

```
ga=1.2; %ga=1.0;  ga=0.7;
```

Levels of membrane voltage

```
U1=-80;
U2=-70;
```

Results of bi-exponential fit of capacitive current

```
A1=MM(1); T1=MM(2); A2=MM(3); T2=MM(4); J01=MM(5); J02=MM(6);
```

Current magnitudes extrapolated to the instant of voltage step

```
J1=A1.*exp(Dt./T1);
J2=A2.*exp(Dt./T2);
```

Access resistance (Eq.10)

```
Ra=(U2-U1)./(J1+J2+J02-J01);
```

Time constant related to surface membrane (Eq 11)

```
Ts=((J1+J2+J02-J01)*T1*T2)/((T1*J2)+(T2*J1));
```

Surface membrane capacitance (Eq 11)

```
Cs=Ts/Ra;
```

Calculation of the resistances $R1 = Rms\ Rt / (Rms+Rt)$ and $R2 = Rms\ (Rms+Rt) / (Rms+Rmt+Rt)$

(Eqs 12-13)

```
b=(T1^2*J2+T2^2*J1)*Ts/((T1.*J2+T2.*J1)*T1*T2);
R1=Ra./(b-1);
a=(J1+J2)/(J1+J2+J02-J01);
R2=Ra*a./(1-a);
```

Calculation of the resistance *R*12 (defined in Eq 18)

```
Re12=Ra*(T1*J2+T2*J1)/(Ts*(J1+J2));
```

Rms and *G*ms=1/*R*ms as a function of variable k (denoted kk) expressed from Eq 17

```
kk=0.05:0.001:1.5;
Rms1=(R1+R2+ga*kk*(R2-R1)+((R1+R2+ga*kk*(R2-R1)).^2-4*R1*R2).^0.5)/2;
Gms1=1./Rms1;
```

*R*ms and *G*ms=1/*R*ms as a function of variable *k* (*kk*) expressed from Eq 18

```
Rms2=((ga*kk-1)*R1*Re12)./(kk*R1-Re12);
Gms2=1./Rms2;
```

Solution of the set of Eqs (17) and (18)

```
q=find(abs((Gms2-Gms1))-min((abs(Gms2-Gms1)))==0);
if q>1
k=kk(q);
end
Gms=Gms2(q(1)); Rms=1/Gms;
```

Grafic solution of the set of Eqs 17 and 18

```
plot(kk,Gms1,kk,Gms2)
```

```
axis([0 0.9 -0.1 0.1])
title('Graphic solution')
xlabel('kk')
ylabel('Gms (uS)')
legend({'Gms1','Gms2'})
grid on
```

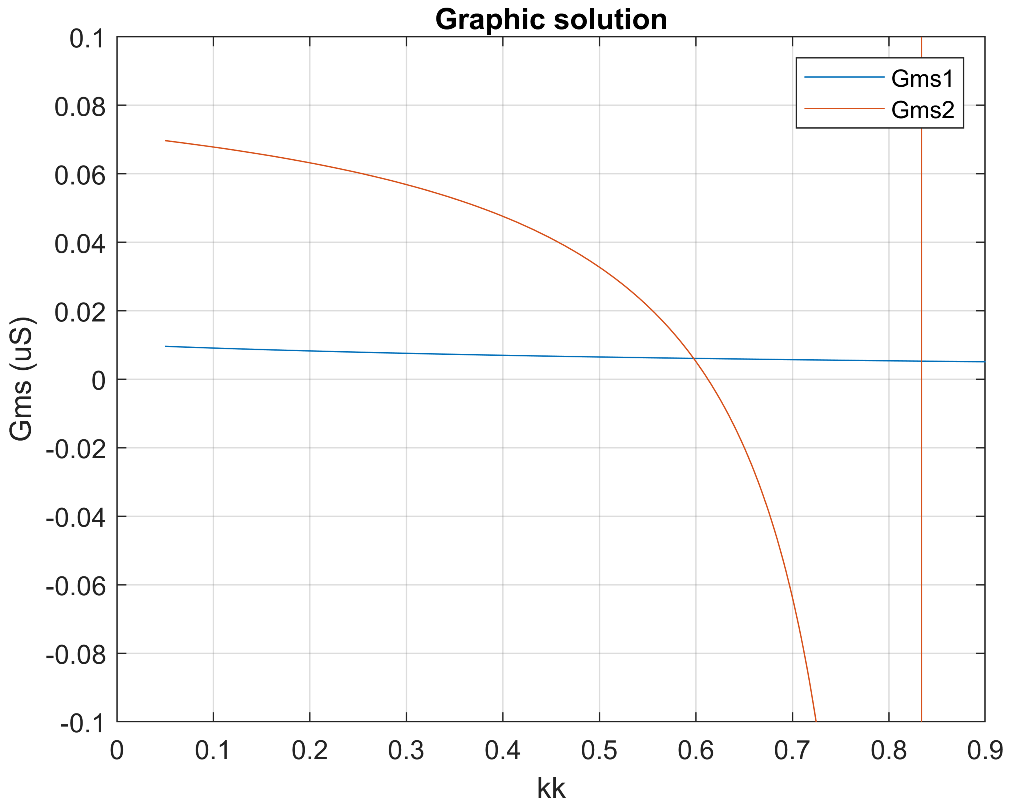


```
% hold on
```

Calculation of *C*t (Eq 19)

```
Ct=k*Cs;
```

Calculation of the resistances *R*t and *R*mt (Eq 20)

```
Rt=R1*Rms/(Rms-1);   Rmt=Rms/(ga*k);
```

Calculation of the total membrane capacitance and the fraction *ft*=S1/(S1+S2) (Eq 21)

```
Cm=Ct+Cs;    ft=k/(k+1);
```

A rough estimate of the resting voltage *U*r, assuming that its value is the same for the surface and tubular membrane; (to check equality, the value of *U*r is calculated from two relations: Ur1 = Ur2).

```
Ur1=U1-J01.*Ra/(1-a);
Ur2=U2-J02.*Ra/(1-a);
```

```
Ur=Ur1;  %Ur=Ur2;
```

Capacitances recalculated from nF to pF

```
Cs=Cs*1000; Ct=Ct*1000; Cm=Cm*1000;
```

Calculated values of the element electrical equivalent circuit (capacitances in pF, resistances in MO, voltage in mV)

```
k
```

```
k = 0.5980
```

```
ga
```

```
ga = 1.2000
```

```
Tab1=table(Cs, Ct, Cm)
```

Tab1 = 1×3 table

| | Cs | Ct | Cm |
|---|---|---|---|
| 1 | 74.2686 | 44.4126 | 118.6812 |

```
Tab2=table(ft, Ra, Rt)
```

Tab2 = 1×3 table

| | ft | Ra | Rt |
|---|---|---|---|
| 1 | 0.3742 | 12.4446 | 14.1040 |

```
Tab3=table(Rms, Rmt, Ur)
```

Tab3 = 1×3 table

| | Rms | Rmt | Ur |
|---|---|---|---|
| 1 | 168.2145 | 234.4126 | -130.0939 |
